# Meaning for reading: The neurocognitive basis of semantic reading impairment after stroke

**DOI:** 10.1101/2025.02.06.636920

**Authors:** Ryan Staples, J. Vivian Dickens, Sara M. Dyslin, Andrew T. DeMarco, Sarah F. Snider, Rhonda B. Friedman, Peter E. Turkeltaub

## Abstract

Diagnosis of alexia has historically focused on syndromes, such as surface alexia, which capture discrete patterns of reading deficits observed in some patients, but do not describe the breadth of reading deficits observed in practice. Aphasia research has recently shifted focus to language process impairments rather than syndromic classifications. A similar shift in focus to reading process impairments may improve our diagnostic approach to alexia. Behavioural evidence suggests that semantic processing influences reading aloud, and that semantic deficits in the context of semantic dementia underlie surface alexia. Because stroke rarely causes loss of semantic knowledge and surface alexia as a syndromic diagnosis is unusual after stroke, semantic reading deficits in stroke alexia have not previously been examined systematically. Semantic reading deficits in stroke may not relate to semantic deficits per se, but rather an inability to use semantics to support reading. Imageability, the degree to which a word brings to mind an image, is one index of semantic influences on reading. High imageability words are read more quickly by healthy persons and more accurately following brain damage. Here, we test if deficits of semantic reading after stroke, as indexed by reduced imageability effects, result from semantic or post-semantic processes. Examining nonverbal semantic processing, semantics-phonology mapping, and semantic control in a sample of 56 left-hemisphere stroke survivors, only semantics-phonology mappings predicted the reading advantage for high imageability words over low imageability words. Support vector regression voxel-based lesion symptom mapping revealed that damage along the superior temporal sulcus and underlying white matter, extending into both the middle and superior temporal gyri, reduced the advantage of high imageability over low imageability words during reading, reflecting an inability to use semantics to support reading. A similar cluster related to impairments in semantics-phonology mapping. The imageability and semantics-phonology mapping results overlapped in the left posterior superior temporal sulcus. Support vector regression connectome lesion symptom mapping revealed white matter disconnections within a broad temporoparietal network important for both phonological and semantic processing were associated with a reduction of the imageability advantage during reading. These results demonstrate that, irrespective of syndromic classification, semantic reading deficits occur in left-hemisphere stroke survivors as a result of impaired integration of semantic and phonological representations, and that the left posterior superior temporal sulcus underlies this process. Our results clarify the neurobiology of reading aloud, and support the existence of a post-semantic impairment of semantic reading in left-hemisphere stroke survivors.

## Introduction

Reading is a cultural invention, learned through years of explicit instruction and effortful practice. Literacy is thus a remarkable achievement. Reading is now so requisite to living in contemporary societies that the loss of fluent reading is a significant cause of disability.^1^ The most common cause of alexia, an acquired reading impairment, is left hemispheric stroke. Alexia usually co-occurs with aphasia, an acquired impairment in language, reflecting the fact that reading is parasitic on language processing abilities.^2^ Current clinical diagnosis relies on alexia syndromes defined by the specific types of words that the person reads aloud inaccurately.^3,4^ Many people with alexia do not fit into a recognized syndrome, and syndromic categorization misses the wide variation in specific deficits within categories. Just as aphasia research and practice has shifted away from syndromes toward process-level descriptions of deficits, examining process-level impairments in alexia irrespective of syndromic classification may reveal previously unrecognized patterns of reading deficits. We take that approach here.

At least two information processing mechanisms enable spelling-to-sound mappings in reading.^5–7^ The first, a sub-word, or sublexical, mechanism decodes sound generatively from print based on direct spelling-to-sound mappings that occur frequently in the language.^6,8^ The second, a word-specific, or lexical, mechanism, ensures correct reading of known words.^6,7^ The two most prominent computational accounts of word reading disagree on the nature of the information processed in the lexical mechanism. The Dual Route Cascaded (DRC) account of reading aloud argues for a reading-specific process: whole word information is represented in parallel orthographic and phonological lexicons, containing lexical units representing whole wordforms with little reliance on semantic content.^6^ In contrast, the primary systems hypothesis argues that reading, as a recent cultural invention, relies on the neural networks that subserve phonology, semantics, and visual processing rather than reading-specific mechanisms.^9^ Connectionist “triangle” models, instantiating the primary systems hypothesis, posit that whole word information is encoded as a distributed semantic representation, the activation of which contributes to pronunciation.^8^

There is significant evidence from computational models, typical readers, and individuals with acquired reading disorders that semantics are recruited to support reading (hereafter, “semantic reading”). Computationally, triangle model implementations demonstrate learning of both direct orthography-to-phonology mappings and indirect orthography-semantics-phonology mappings.^7^ Lesioning the semantic route of these models causes a form of reading impairment called surface alexia^7,10,11^, characterized by reduced accuracy reading low-frequency irregular words, for which print-to-sound mappings are statistically improbable^3^, and the production of regularization errors, where irregular words are pronounced as if they were regular (e.g., *pint* [/ pa nt/] pronounced to rhyme with *mint* [/ m nt/]).^3^

Surface alexia is common in people with loss of semantic knowledge due to semantic dementia (SD).^11,12^ SD is a neurodegenerative condition that causes progressive anterior-to-posterior atrophying of the lateral temporal lobes and loss of semantic knowledge, reflected in poor performance on tasks such as picture naming or matching pictures based on semantic similarity.^13,14^ The degree of semantic impairment in SD and the emergence of surface alexia are correlated.^11^ Triangle model accounts of reading draw on the associations between SD, impaired semantic cognition, and surface alexia to argue that the anterior temporal lobes, particularly in the left hemisphere, support semantic reading through the storage or processing of semantic representations. The DRC account holds that semantics are not required for reading, and that there should be no principled relationship between semantic impairment and reading aloud.^6,15^ Per the DRC account, the relationship between semantic impairment in SD and surface alexia is due to the spatial proximity of cortical regions that separately support each function: damage to the anterior temporal lobes impairs semantic cognition, and additional atrophy in posterior regions, such as the ventral occipitotemporal cortex, impairs word reading.^15^

Reflecting the distributed nature of semantic representation in the brain, loss of semantic knowledge is rare in patients with focal brain injuries, including stroke. While less common after stroke relative to SD, symptoms of surface alexia – regularization error production and poor reading of low-frequency irregular words – do sometimes occur after stroke. For example, a prior lesion-mapping study found that regularization error production was associated with damage to the posterior temporal white matter and MTG.^16^ The dissociation between intact semantic knowledge and surface alexia symptoms in stroke might suggest that surface reading deficits in stroke can arise due to non-semantic lexical reading deficits, as proposed in the DRC model. Alternatively, surface reading deficits might reflect an inability to apply intact semantic knowledge to reading aloud, due to deficits in mapping semantics to phonology or to impaired executive control of semantics, which has been suggested to underlie semantic deficits in stroke aphasia.^17,18^ Because surface reading deficits can be explained theoretically without the involvement of semantics in reading aloud, the occurrence of these deficits alone does not necessarily indicate a deficit in semantic reading. Thus, a more direct measure of semantic processing for reading aloud is needed to clarify the nature of semantic reading deficits after stroke.

A critical line of behavioral research supporting the importance of semantics for reading aloud concerns effects of word concreteness or imageability. Imageability is a measure of the degree to which a word’s referent evokes a mental image, such that ‘*dog*’ is highly imageable whereas ‘*honor*’ is not.^19^ Imageable words are read faster and more accurately by healthy neurotypical readers.^19,20^ This imageability effect is thought to occur because imageable words have more robust semantic representations^21^, perhaps due to embodied, perceptual features not shared by low imageability words.^22^ In stroke alexia, imageability or concreteness effects are often considered a sign of sublexical (or phonological) reading deficits. Individuals with severe sublexical reading deficits sometimes show exaggerated imageability effects, thought to reflect a reliance on semantic reading when direct spelling-to-sound translation is impaired.^23^ In contrast, a loss of the advantage for high imageability words over low imageability words would provide direct evidence for a semantic reading deficit.

The brain regions required for an intact imageability advantage are unclear. fMRI studies of reading aloud^24^ and lexical decision^25,26^ in typical adults suggest that the bilateral angular and posterior cingulate gyri are activated preferentially for high as opposed to low imageability words. Individual differences in typical adults’ white matter volume in tracts connecting the left inferior temporal sulcus, posterior middle temporal gyrus, and angular gyrus are related to variation in the imageability effect.^27^ In the only lesion study to date on concreteness or imageability effects in reading, Dickens et al.^28^ identified a region spanning the pars orbitalis and pars triangularis of the left inferior frontal gyrus (lIFG) in which lesions reduced the reading advantage for concrete words relative to abstract words. The lIFG is a critical region for semantic control^17^, suggesting that semantic contributions to reading aloud might rely on the capacity for flexible use of semantic information. The interpretability of this result was limited, however, as the stimulus set did not induce a group-level concreteness effect, and the relationship between reading deficits and semantic impairments was not asssessed. In sum, this work implicates a network of left posterior temporoparietal and inferior frontal regions in semantic reading. Whether lesions to specific regions or connections in this network produce semantic reading deficits remains untested.

Presently, we examine the imageability effect in reading aloud after left hemisphere stroke to clarify the nature of any semantic reading deficits in stroke alexia, specifically examining their relationships with nonverbal semantics, semantic-to-phonology (S-P) mapping, and semantic control. We then perform voxel-based (VLSM) and connectome lesion-symptom mapping (CLSM) to identify lesions that result in semantic reading deficits as reflected by a reduced imageability effect. The results provide direct evidence of semantic reading deficits after stroke that are related to impairments in S-P mapping, and are associated with lesions to the left STS and connections within a broader temporoparietal semantic network.

## Materials and methods

### Participants

Participants included 56 people with left-hemisphere stroke and 68 age– and education-matched neurotypical controls (Table 1; see Supplementary information for recruitment details). All participants in the stroke cohort were at least six months post-stroke at the time of enrollment. Written informed consent was obtained from all participants as required by the Declaration of Helsinki. The study protocol was approved by the Georgetown University Institutional Review Board.

**Table 1.**
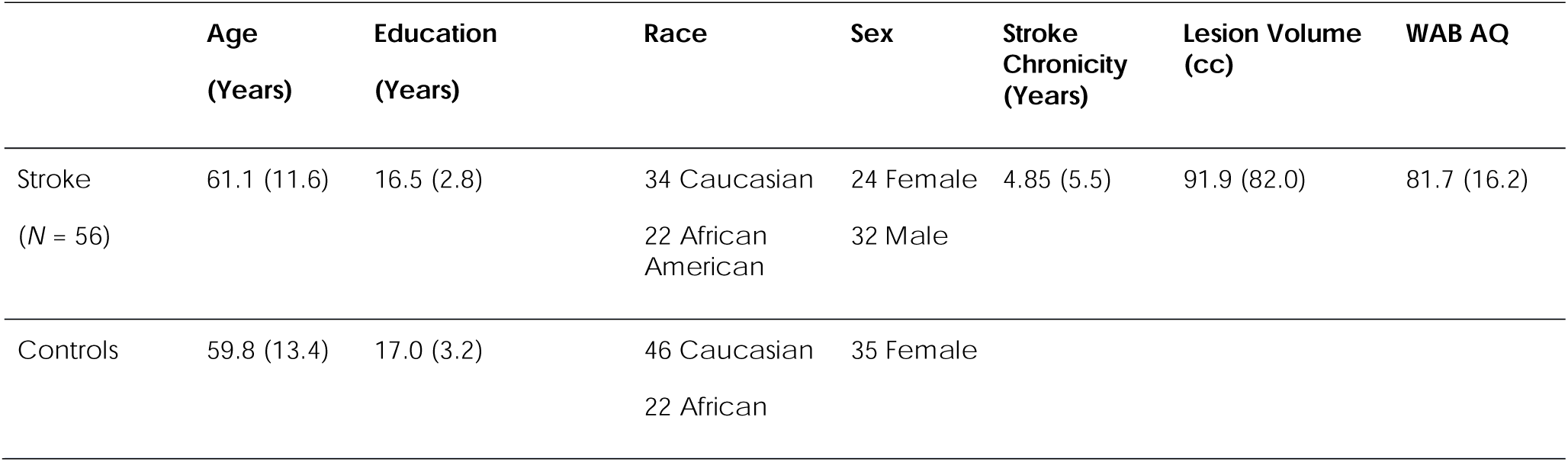
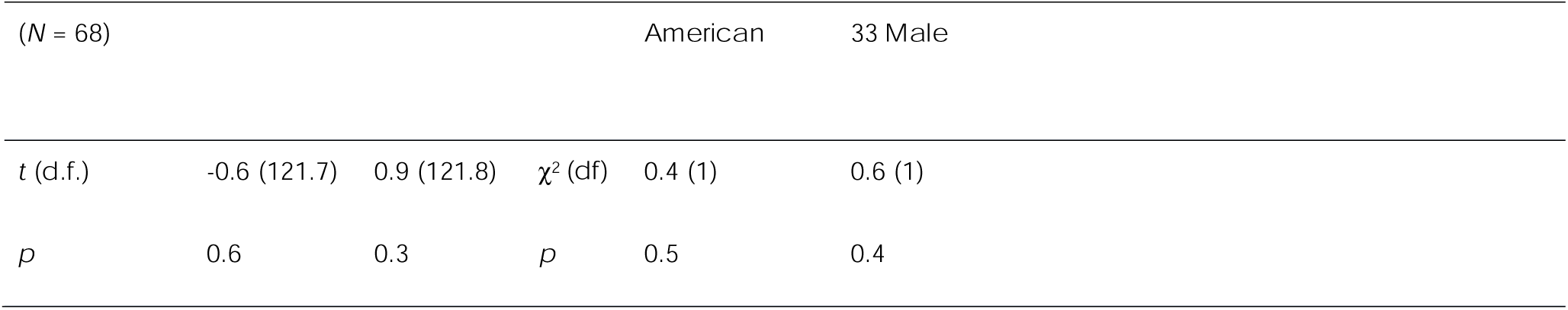
Participant demographics and stroke participant clinical data.

### Cognitive tests

#### Oral reading test

Participants read aloud a list of 200 monosyllabic English words that varied categorically in word frequency (low versus high frequency of occurrence in English, such as *sheen* versus *lunch*), spelling-to-sound regularity (regular versus irregular spelling-to-sound correspondences, such as *steam* vs *steak*), and imageability^19,20^ (low versus high mental imagery, such as *grace* versus *horse*). The word list consisted of 100 words within each level of each factor. Letter length ranged from three to six letters. Low and high imageability words were matched on letter length, frequency, regularity, and articulatory complexity.^29^ Matching between word types was assessed through Welch’s two-sample t-tests. See Supplementary material for additional information on word selection, matching, the item list and associated psycholinguistic features, as well as test administration and scoring details. Briefly, each word was displayed in a pseudorandomized order in the center of a touchscreen. Participants were instructed to read aloud the word as quickly and accurately as possible within a ten-second limit. Accuracy was scored for the first complete attempt, defined as the first response consisting of a consonant and vowel (CV or VC), excluding a schwa /ə/.^30^ An isolated vowel was accepted as the first complete attempt if the target itself was a CV or VC.

#### Semantic tests

The stroke cohort completed non-orthographic language tasks designed to assess 1) non-verbal semantics, 2) the ability to map semantics to phonology (S-P mapping), and 3) the controlled use of semantic processing (semantic control). Nonverbal semantics was measured by averaging the accuracies on two tests of associative semantic knowledge: Pyramids and Palm Trees^31^ and the Temple Assessment of Language and Short-term Memory in Aphasia (TALSA) category judgment subtest.^32^ For Pyramids and Palm Trees, the subject was presented a picture on a touchscreen (e.g., a pyramid) and matched it to one of two pictures presented below: the semantically related target (e.g., a palm tree) or the incorrect distractor (e.g., a pine tree; 52 trials). The TALSA category judgment subtest measured knowledge of animals, transportation, vegetables, furniture, and musical instruments through 60 pairs of pictures. Two pictures were displayed in succession for three seconds each (e.g., banana and apple), and the subject had to indicate whether the two pictures belonged to the same category by pressing “yes” or “no” on the touchscreen.

S-P mapping ability was measured by averaging accuracy on an in-house auditory word-to-picture matching test (48 trials) and on two picture naming tests (the 60-item Philadelphia Naming Test^30^ and an in-house 60-item picture naming test).^33^ For auditory word-to-picture matching, the participant was presented a word auditorily via headphones and selected the corresponding picture among five semantically related foils on a touchscreen. For the picture naming tests, participants were instructed to use one word to name each picture as quickly and accurately as possible, within a 20-second limit. Accuracy was scored on the first complete attempt. Combining auditory word-to-picture matching and picture naming produced a measure of bidirectional semantic-phonological processing (but see Supplementary Tables 3-6 and Supplementary Figures 1-2 for separate analyses of picture naming and auditory word-to-picture matching).

Semantic control was measured using a previously validated four-alternative forced choice task.^34^ Participants were instructed to touch a target item on a touchscreen as many times as possible within 20-second blocks. The target item changed location after each touch. Participants had to tap either 1-, 2-, or 3-item sequences, depending on the block. To assess semantic control, two versions of the task were contrasted: a semantic version, in which the four items were semantically related, and an unrelated version, in which the four items were neither semantically nor phonologically related. A Time Per Target Selection (TTS) score was calculated for each version (semantic and unrelated) by dividing the block length (20 seconds) by the number of times a participant selected the target(s) in each block, averaging across the blocks, and then taking the natural log of the resulting value. The TTS on the unrelated version was subtracted from the semantic version score to determine a participant-level semantic control score.^34^

### Neuroimaging

#### MRI acquisition and lesion tracing

The following sequences were acquired with Georgetown’s 3T Siemens MAGNETOM Prisma scanner using a 20-channel head coil, as previously described in Dickens et al.^35^: a T1-weighted magnetization prepared rapid gradient echo (MPRAGE) sequence (1 mm^3^ voxels), a fluid-attenuated inversion recovery (FLAIR) sequence (1 mm^3^ voxels), and a high angular resolution diffusion imaging (HARDI) sequence (81 directions at *b* = 3000, 40 at *b* = 1200, 7 at *b* = 0; 2 mm^3^ voxels). For stroke subjects, author P.E.T. manually traced stroke lesions on the native-space MPRAGE and FLAIR images via ITK-SNAP.^36;^ http://www.itksnap.org/ MPRAGEs and lesion tracings were warped to the Clinical Toolbox Older Adult Template^37^ using Advanced Normalization Tools^38^ as described in Dickens et al.^28^. See the Supplemental Materials for additional neuroimaging acquisition details.

#### Structural connectome pipeline

MRtrix 3.0^39^ software was used to preprocess the HARDI data, estimate white matter pathways, and create a structural connectome for each subject. Multi-shell, multi-tissue constrained spherical deconvolution^40^ was applied to calculate the voxelwise fiber orientation distributions from the preprocessed HARDI data. Probabilistic anatomically constrained tractography^41^ on the white matter fiber orientation distributions traced 15 million streamlines in native space (algorithm = iFOD2, step = 1, min/max length = 10/300, angle = 45, backtracking allowed, dynamic seeding, streamlines cropped at grey matter-white matter interface). The Spherical Deconvolution Informed Filtering of Tractograms-2 algorithm^42^ was applied to derive cross-sectional multipliers for each streamline to adjust streamline densities to be proportional to the underlying white matter fiber densities. For each subject, a white matter connectivity matrix was created by assigning streamlines to brain parcels of the Lausanne atlas at scale 125 (https://github.com/mattcieslak/easy_lausanne)^43^. Overall, each edge (connection) of a connectome represents the apparent fiber density of the white matter connecting the two brain parcels. For the purpose of structural connectome lesion-symptom mapping, each stroke subject’s connectome was binarized by comparing stroke and control connectomes to label each connection as lesioned (0) or intact (1). Specifically, a connection in a stroke subject’s connectome was deemed lesioned if the apparent fiber density was below the first percentile of the control values for that specific connection. See the Supplemental Materials for additional details on brain parcellation and connectome construction.

### Statistical Analyses

#### Behavioral analyses

We estimated two multiple linear regression models to determine if the three measures of semantic cognition (nonverbal semantics score, S-P mapping score, and semantic control score) were related to semantic reading ability. The dependent variables of the two regressions were 1) accuracy on high imageability words and 2) accuracy on low imageability words. Predictors in both regressions were the non-verbal semantics score, the S-P mapping score, and the semantic control score, covarying for accuracy on low imageability words in the first model and for accuracy on high imageability words in the second model. Additional covariates included in each regression were age, education, and lesion volume.

#### Voxel-based lesion-symptom mapping

We conducted two support-vector regression VLSM (SVR-VLSM)^44^ analyses to identify brain regions important for semantic contributions to reading aloud by examining the relationship between lesion location and accuracy reading high imageability words; the first analysis examined all high imageability words, and the second examined high imageability irregular words because the triangle model predicts that semantic contributions to reading are particularly important for irregular words.^10^ Both analyses covaried for accuracy on low imageability words (all low imageability words in the first analysis, and low imageability irregular words in the second) to control for the contribution of non-semantic processes to reading accuracy, including visual processing, direct orthography-to-phonology mapping, and articulation. Additional covariates included lesion volume, age, and education, which were regressed out of both the lesion and behavioural data prior to SVR.^45^

To identify brain regions required for intact semantic cognition, we conducted SVR-VLSM analyses of 1) the nonverbal semantics score, 2) the S-P mapping score, and 3) the semantic control score. Covariates regressed out of the behavioral and lesion data included the two semantic variables that were not the target of the current analysis, lesion volume, age, education, and accuracy reading low imageability words. These analyses covaried accuracy reading low imageability words to control for any contribution of phonological processing deficits to accuracy on the semantic tests (e.g, phonological working memory).

For all SVR-VLSM analyses, only voxels lesioned in at least 10% of stroke participants (i.e., at least 6) were included. Statistically significant lesion-behavior relationships were determined through permutation testing. Behavioral scores were randomly reassigned to subjects over 10,000 permutations, and a distribution of SVR β-values was calculated on a voxel-wise basis (*P* < 0.005, one-tailed (negative)).^46^ Cluster-wise significance was then determined by calculating the cluster size threshold at a family-wise error rate correction of *P* < 0.05. All SVR-VLSM analyses were implemented in the SVR-LSM MATLAB toolbox (https://github.com/atdemarco/svrlsmgui/).^45^

#### Connectome lesion-symptom mapping

We conducted five CLSM analyses matching the VLSM analyses described above (all high imageability words, high imageability irregular words, nonverbal semantics score, S-P mapping score, semantic control score), which identified disconnections between brain parcels associated with reduced scores on the dependent variable. The logic and covariates of these analyses are as described for the VLSM analyses above.

Only connections present in 100% of control subjects were included in the SVR to reduce Type I error due to erroneous streamlines. The SVR-CLSM analysis also excluded all intrahemispheric connections within the right hemisphere, given that we examined a group of left hemisphere stroke subjects. Left-hemispheric and inter-hemispheric connections lesioned in at least 10% of the stroke subjects were included in the SVR. Edge-wise (disconnection between two brain parcels) and parcel-wise (disconnection of a single brain parcel, including all anatomical endpoints) significance was calculated via permutation testing (10,000 permutations, CFWER control with v=10, FWER=0.05, one-tailed (negative)).^47,48^

## Data availability

Data may be made available upon request.

## Results

### Summary of behavioral performance

Stroke survivors read aloud all word types less accurately than healthy controls (Table 2). The stroke cohort demonstrated a wide range in reading accuracy, as expected given the cohort’s variability in stroke size and aphasia severity. Stroke participants read high imageability words aloud more accurately than low imageability words (stroke survivor high imageability advantage=6.7%). Further, the stroke survivors overall showed an enhanced imageability effect compared to the controls (control mean high imageability advantage=2.5%, *t*(59.6)=-4.04, *p*<0.001). That is, low imageability word reading is affected more by stroke in general, while high imageability words are relatively spared, consistent an overall reliance on semantics for reading aloud after stroke. The high imageability advantage in both stroke survivors and controls was largest, on average, for low frequency words (imageability advantage=9.0% in stroke survivors and 4.2% in controls) and irregular words (imageability advantage=10.4% in stroke survivors and 5.6% in controls).

**Table 2.** Summary of participant performance while reading aloud.

|  |  | HI | LI | HI LF | LI LF | HI HF | LI HF | HI Irr | LI Irr | HI Reg | LI Reg |
| --- | --- | --- | --- | --- | --- | --- | --- | --- | --- | --- | --- |
| Healthy Controls | Mean (SD) | 96.9 (4.3) | 94.4 (5.9) | 94.9 (6.8) | 90.7 (9.3) | 98.8 (2.2) | 98.0 (3.0) | 95.3 (5.3) | 89.7 (8.9) | 98.4 (3.9) | 99.0 (3.2) |
|  | Range | 74 - 100 | 68 - 100 | 58 - 100 | 52 - 100 | 86 - 100 | 84 - 100 | 72 - 100 | 54 - 100 | 76 - 100 | 82 - 100 |
| Stroke Survivors | Mean (SD) | 80.2 (21.3) | 73.6 (24.8) | 74.6 (24.2) | 65.6 (27.1) | 85.9 (19.1) | 81.5 (23.8) | 77.6 (20.5) | 67.3 (24.0) | 82.9 (23.0) | 79.9 (26.4) |
|  | Range | 16 - 100 | 3 - 98 | 10 - 100 | 4 - 96 | 22 - 100 | 2 - 100 | 16 - 100 | 4 - 98 | 16 - 100 | 2 - 100 |
|  | *t* (d.f.) | 5.8 (56.9) | 6.2 (57.6) | 6.2 (58.7) | 6.8 (60.4) | 5.1 (55.6) | 5.2 (55.7) | 6.4 (58.1) | 6.8 (61.5) | 5.0 (56.3) | 5.4 (55.7) |
|  | *p* | <0.001 | <0.001 | <0.001 | <0.001 | <0.001 | <0.001 | <0.001 | <0.001 | <0.001 | <0.001 |

HI = high imageability. LI = low imageability. HF = high frequency. LF = low frequency. Irr = irregular. Reg = Regular.

Stroke survivors’ performance on the semantic composites is shown in Table 3. Consistent with the tendency of stroke survivors with aphasia to have greater phonological as compared to semantic deficits, nonverbal semantic task performance was good whereas SP task performance was more impaired. Semantic control values were within the range of healthy controls.^34^

**Table 3.** Summary of participant performance on semantic tasks.

|  | S-P Mapping Score | Nonverbal Semantics Score | Semantic Control Score |
| --- | --- | --- | --- |
| Mean (SD) | 80.7 (17.4) | 91.0 (7.2) | -0.4 (0.2) |
| Range | 31.0 - 98.9 | 65.0 - 99.1 | -1.2 - 0.0 |

### Relationships between imageable word reading and semantic processing

Two independent regression models assessed the relationship between semantic processing and reading accuracy on high or low imageability words. Reading accuracy on high imageability words was predicted by accuracy on low imageability words (*p*<0.001) and by the S-P mapping score (*p*=0.046), but not by nonverbal semantics or semantic control (*p*’s>0.05). Accuracy reading low imageability words was predicted by accuracy reading high imageability words (*p*<0.001), but not by nonverbal semantics, S-P mapping, or semantic control (*p*’s>0.05) (Table 4). These analyses demonstrate that the advantage for high imageability words in oral reading relates to semantic-to-phonological mapping. Next, we examined whether and how lesions to specific brain structures reduce this reliance on semantics during oral word reading.

**Table 4.**
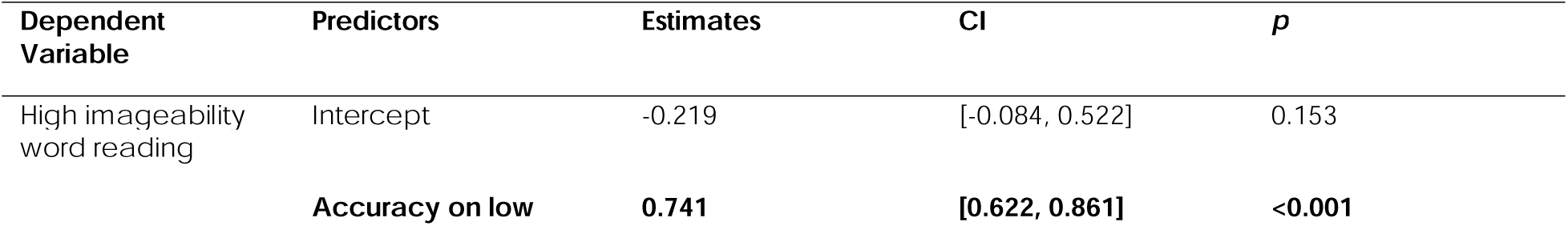
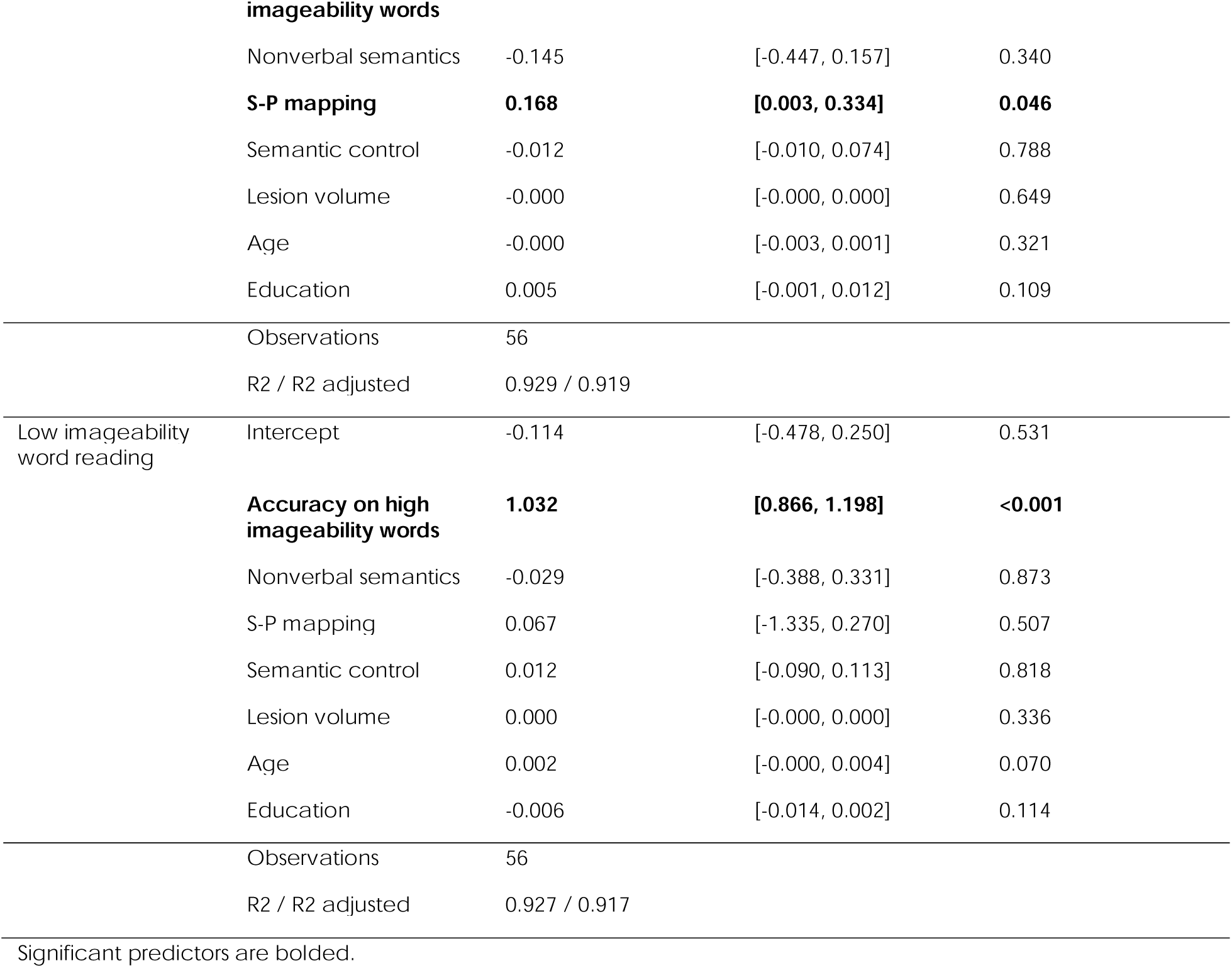
Model predicting high imageability word reading accuracy.

### Lesion-symptom mapping

The study samples’ lesion distribution was typical of left hemisphere stroke cohorts with aphasia, with most lesions being within the left middle cerebral artery (MCA) territory (Fig. 1). SVR-VLSM revealed a significant association between posterior left superior temporal sulcus (STS) lesions (MNI centroid: –54.9, –32.2, –5.0) and lower accuracy on high imageability words, controlling for accuracy on low imageability words (Table 5; Fig. 2A). In other words, lesions of the left posterior STS reduce the typical advantage of high imageability in oral word reading. The association between left STS lesions remained significant when restricted to high imageability irregular words only (MNI centroid: –51.4, –36.0, 2.4). This result overlapped with the all-high-imageability-word result in the STS and underlying temporal white matter (MNI centroid: –52.5, –35.0, 1.5; all high imageability words: 5.2 cc; high imageability irregular words: 4.8cc; overlap: 2.9cc; Dice coefficient: 0.58) (Table 5, Fig. 2B).

**Fig. 1.**
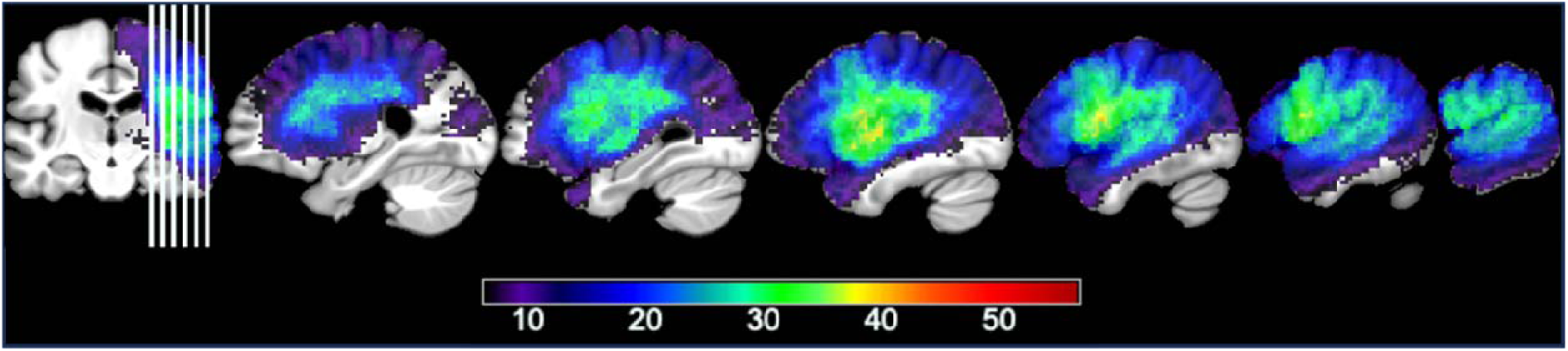
– Voxelwise lesion overlap map.

**Fig. 2.**
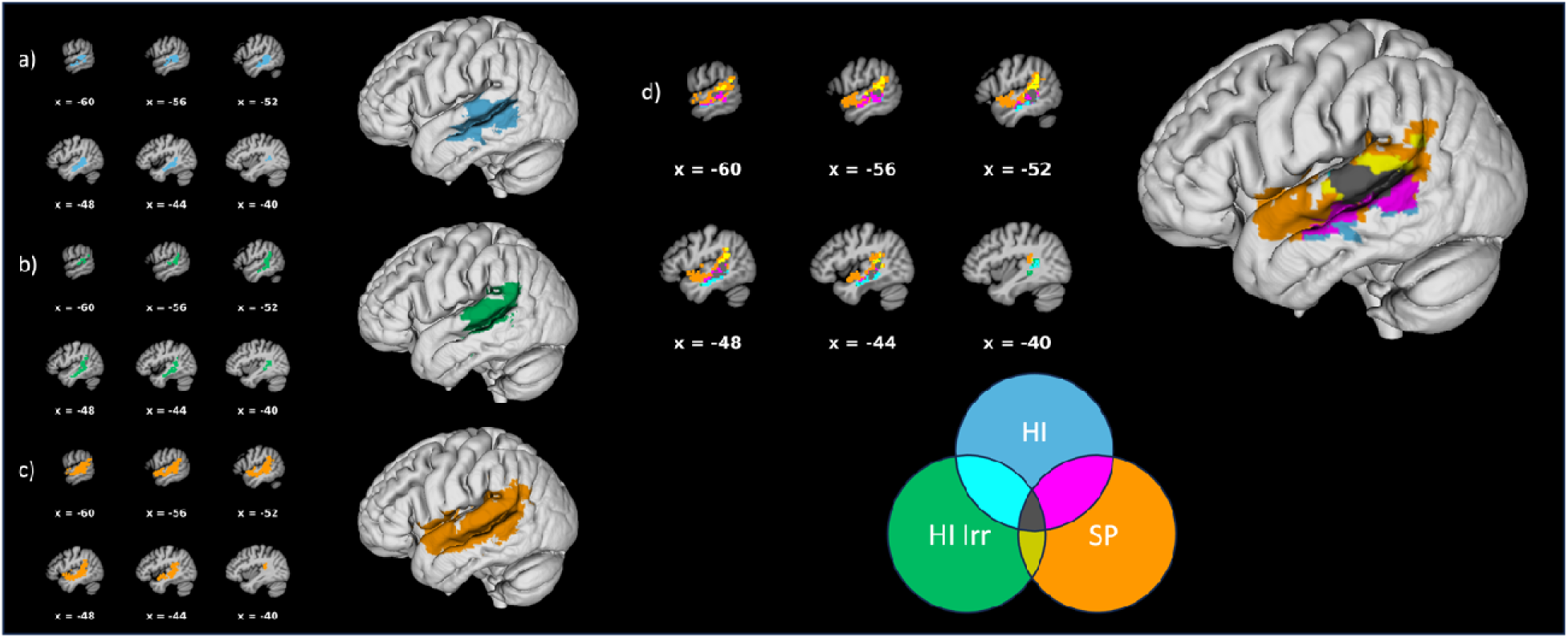
– SVM-VLSM results at voxelwise *p* < 0.005 and clusterwise FWER *p* < 0.05. a) Reduced accuracy on high imageability word reading relative to low imageability word reading (clusterwise *p* = 0.031). b) Reduced accuracy on high imageability, irregular word reading relative to low imageability, irregular word reading (clusterwise *p* = 0.040). c) Reduced S-P scores, controlling for nonverbal semantic and semantic control scores (clusterwise *p* = 0.0028). d) Overlap of all high imageability word reading accuracy, high imageability irregular word reading accuracy, and S-P score SVR-VLSM results. Age, education, and lesion volume are regressed out of all analyses.

**Table 5.**
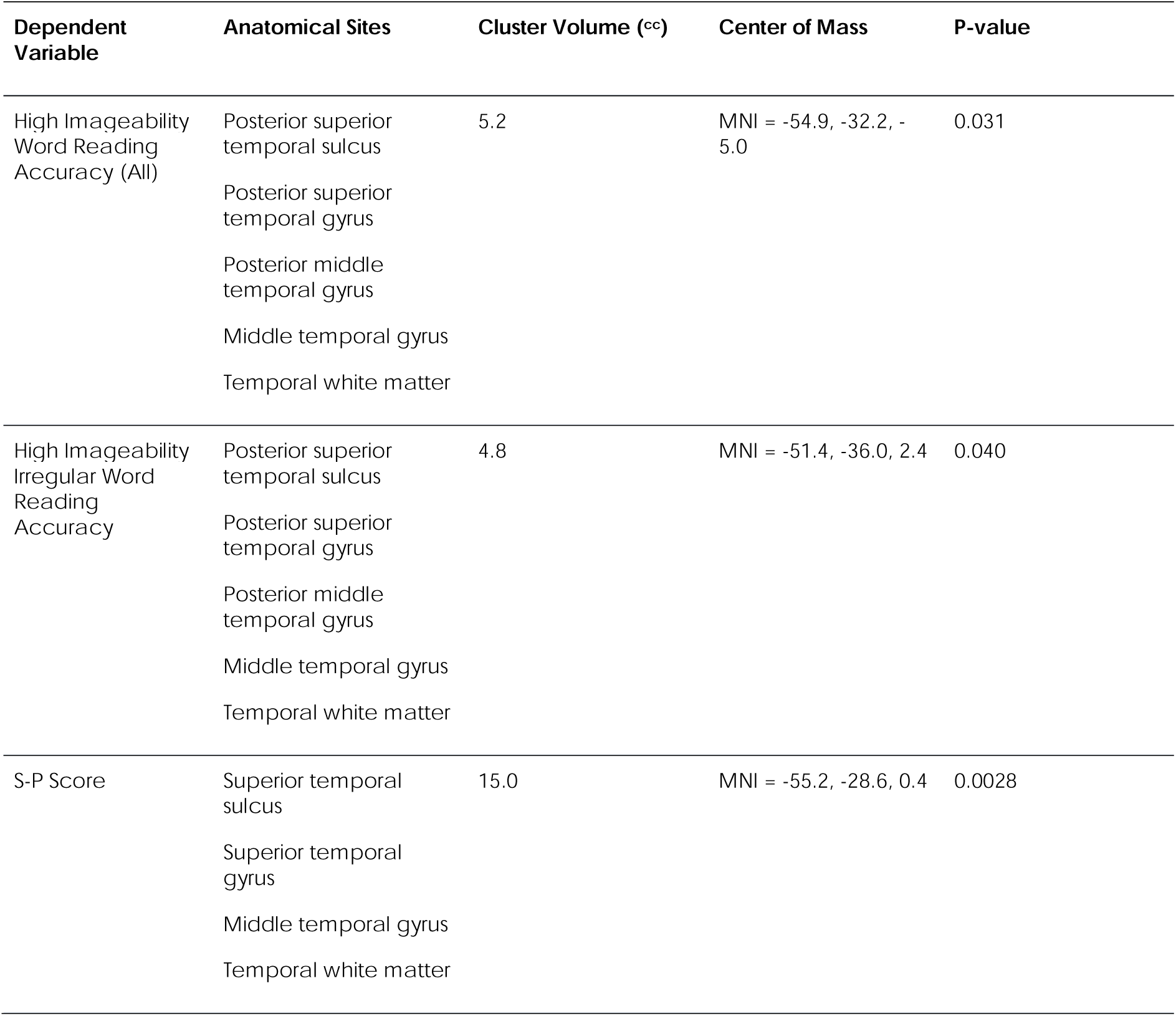
VLSM-SVM results (Cluster FWER *P* < 0.05)

| Dependent Variable | Anatomical Sites | Cluster Volume (cc) | Center of Mass | P-value |
| --- | --- | --- | --- | --- |
| High Imageability Word Reading Accuracy (All) | Posterior superior temporal sulcus | 5.2 | MNI = -54.9, -32.2, -5.0 | 0.031 |
|  | Posterior superior temporal gyrus |  |  |  |
|  | Posterior middle temporal gyrus |  |  |  |
|  | Middle temporal gyrus |  |  |  |
|  | Temporal white matter |  |  |  |
| High Imageability Irregular Word Reading Accuracy | Posterior superior temporal sulcus | 4.8 | MNI = -51.4, -36.0, 2.4 | 0.040 |
|  | Posterior superior temporal gyrus |  |  |  |
|  | Posterior middle temporal gyrus |  |  |  |
|  | Middle temporal gyrus |  |  |  |
|  | Temporal white matter |  |  |  |
| S-P Score | Superior temporal sulcus | 15.0 | MNI = -55.2, -28.6, 0.4 | 0.0028 |
|  | Superior temporal gyrus |  |  |  |
|  | Middle temporal gyrus |  |  |  |
|  | Temporal white matter |  |  |  |

Additional VLSM analyses localized the effects of the three semantic variables. No lesions related to nonverbal semantic processing or semantic control. However, lower S-P score were associated with damage along the STS and underlying white matter (MNI centroid: –55.2, – 28.6, 0.4, cluster size: 15.0cc, Table 5, Fig. 2C). This cluster overlapped with the SVR-VLSM result for high imageability words (MNI centroid: –55.6, –36.1, –1.0, overlap: 3.5cc; Dice coefficient: 0.35) and for high imageability irregular words (MNI centroid: –57.8, –35.5, 4.7, Dice coefficient: 0.33). All three clusters overlapped in the dorsal bank of the posterior STS (MNI centroid: –57.5, –35.0, 4.0; overlap cluster size: 1.2cc; Dice coefficient: 0.11, Fig. 2D).

### Connectome lesion-symptom mapping

SVR-CLSM analyses associated lower accuracy on all high imageability words, controlling for low imageability word accuracy with 10 disconnections in a left-lateralized temporo-parieto-occipital network. This network largely contained areas in the semantic network^49^, including the anterior temporal lobe, angular gyrus, and connections throughout the inferior, middle, and superior temporal gryi (Supplementary Table 5, Fig. 3A).

**Fig. 3.**
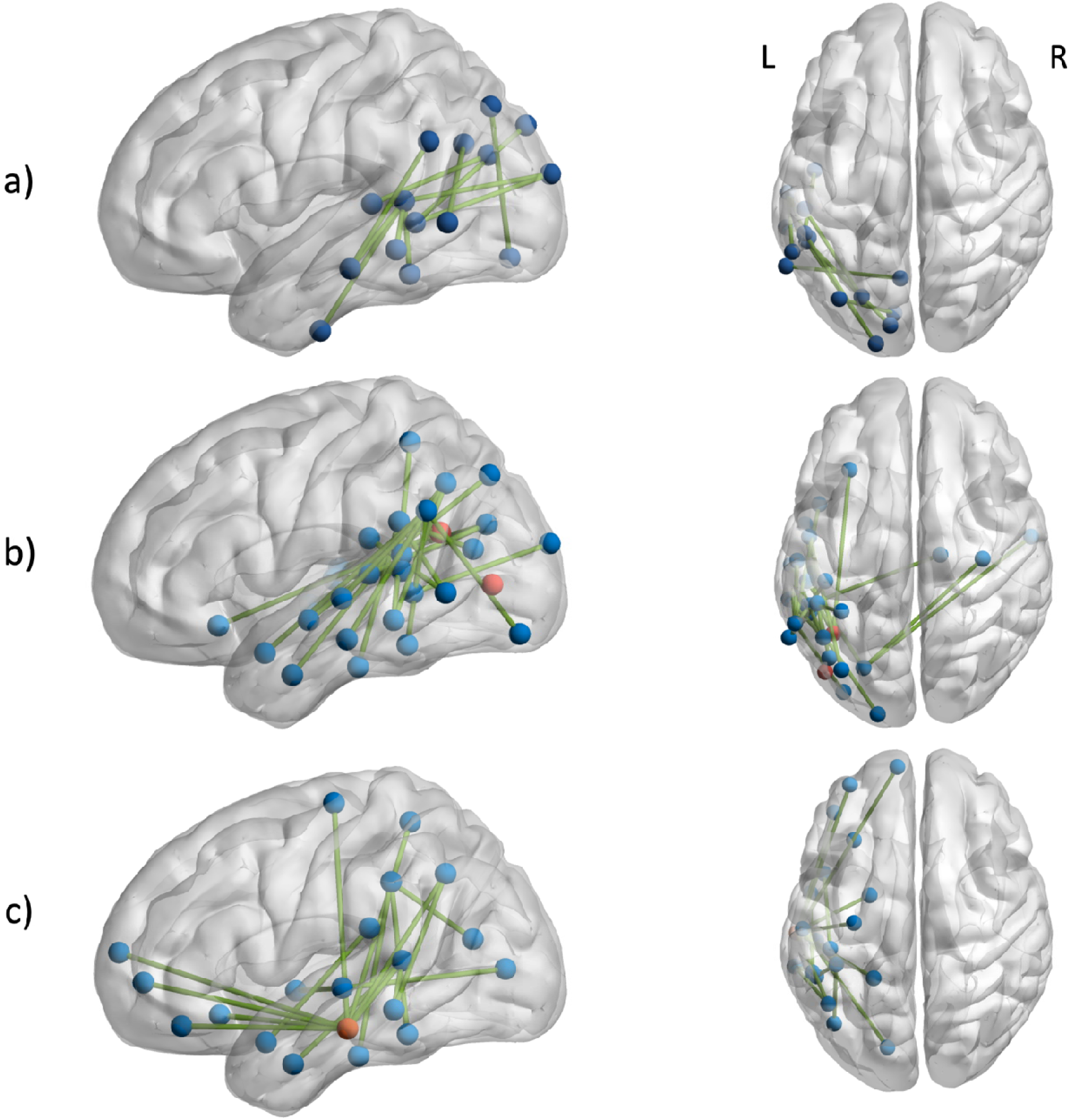
– SVR-CLSM results (edgewise and parcelwise FWER *p* < 0.05) showing. a) edges where disconnections are associated with reduced high imageability word reading accuracy (controlling for low imageability word reading accuracy), b) edges where disconnections are associated with reduced high imageability irregular words (controlling for low imageability irregular words), and c) edges where disconnections related to reduced S-P mapping scores. Orange indicates nodes significant at the parcel level. SVR-CLSM results visualized with BrainNet Viewer ^50^.

Deficits in reading high imageability irregular words, controlling for low imageability irregular words, were associated with denser but spatially similar left-lateralized disconnections. Specifically, 22 disconnections were identified. Sixteen of these disconnections were between the angular gyrus or supramarginal gyrus and parcels spanning the temporal lobe. An angular gyrus parcel (MNI coordinates: –39, –56, 23) and a lateral occipital cortex (LOC) parcel (MNI coordinates: –43, –75, 4) were significant at the parcel level (parcel FWE corrected *P*<0.05). (Supplementary Table 5, Fig. 3B).

The S-P mapping score SVR-CLSM analysis associated lower S-P scores with 17 disconnections (Supplementary Table 5, Fig. 3C). Like the networks associated with the reading of imageable words, most disconnections associated with lower S-P score were between parcels in the temporal and parietal lobes. Eight of the 17 disconnections included a middle temporal gyrus parcel, which itself was significant at the parcel level (MNI coordinates: –60, –22, –26). Additionally, disconnections spanning the inferior, middle, and superior frontal gyrus and this MTG parcel were associated with lower S-P scores.

## Discussion

We aimed to examine the neurocognitive bases of semantic reading impairment in stroke alexia. Using the imageability effect as a direct measure of semantic influence on reading, we find a generally increased reliance on intact semantic processing for reading in stroke survivors compared with typical controls, likely reflecting the frequent impairment of phonological reading mechanisms after stroke.^2^ This demonstrates that for many stroke survivors, reading success depends in part on their ability to use semantics to support reading. The reading deficits observed for words without vivid meanings (low imageability) are thus partially rescued for words with vivid meanings (high imageability). Leveraging variability within the stroke survivors’ imageability effect, we found robust evidence that inability to utilize semantics for oral reading is associated with left posterior temporal lesions and disconnections between posterior temporal and inferior parietal regions of the semantic network. Finally, we found that S-P mapping, but not nonverbal semantic processing or semantic control, predicted accuracy on high imageability word reading. Overall, these findings reveal a post-semantic reading deficit that occurs in stroke alexia, differing from previously described semantic reading deficits associated with surface alexia in SD.

### Neural substrates of semantic influences on reading

Our VLSM findings indicate that the posterior temporal cortex is essential for producing the imageability advantage when reading aloud. Previous work suggests that phonological wordforms are represented by connections between phonological processes in the STG^51^ and the pSTS^52^, with the pSTS directly implicated in retrieving phonological codes.^53,54^ The pSTS also has anatomical connections to anterior parts of the ventral occipitotemporal (vOT) cortex^55^, which responds to printed whole wordforms.^56^ This connectivity profile makes the pSTS a strong candidate as a convergence zone for phonological and orthographic information.^52^ Our results further suggest that the pSTS is involved in processing semantic contributions to reading aloud, as indexed by the imageability effect.

Supporting the role of the pSTS in semantic reading, many of the disconnections identified in the CLSM analyses of high imageability words comprise regions in the semantic network. The semantic network involves a bilateral, but left-biased network of frontal, temporal, and parietal cortex.^49,57,58^ The angular gyrus (AG) is a node in the semantic network^49^, thought to serve as a hub connecting language, experiential representation, and semantic control networks.^59^ Previous evidence links greater AG connectivity to the imageability effect in healthy adult readers.^27,60^ In the VLSM of high imageability irregular words, we identified an AG parcel (MNI coordinates = –39, –56, 23) for which anatomical disconnections overall were associated with a reduced imageability advantage, confirming the role of intact connectivity to an important semantic hub in reading aloud.^17^ The middle temporal gyrus (MTG) was also implicated in the imageability analyses. Lesions to the MTG impair single word comprehension.^61,62^ The posterior MTG has been linked to both phonological wordform-meaning mapping^62,63^ and to the flexible use of semantics in response to task demands.^64,65^ While an impairment of either process could plausibly impair semantic contributions to reading, semantic control was not related to behavior, lesions, or disconnections in our sample. Instead, lesions to the MTG were related to S-P mappings, supporting the wordform-meaning mapping interpretation.

Evidence from SD suggests that the left anterior temporal lobe is a hub of semantic processing.^13,17,66–69^ Both CLSM analyses identified posterior lATL disconnections that were associated with reductions of the imageability advantage. This is consonant with evidence that the imageability effect in reading aloud is sometimes reversed in semantic dementia.^70^ Our lesion coverage of the ventral ATL was limited, so the lack of VLSM findings should not be over-interpreted. The CLSM results suggest that the ATL may contribute to use of semantic knowledge for reading, but future research should investigate if direct damage to the ATL is sufficient to alter the imageability effect.

CLSM also identified disconnections to regions of the brain linked to phonological processing in reading, including the supramarginal gyrus (SMG).^71–73^ Damage to the SMG can cause conduction aphasia^74^ and impairments in rhyme processing.^75^ Disconnections to the superior temporal gyrus (STG) were also associated with reductions of the imageability effect, especially for irregular words. The STG is also involved in phonological processing, both in speech and reading.^51,76,77^ The STG disconnections we identify are primarily to nodes in the semantic network, particularly the AG, highlighting that the imageability effect arises in reading through interactions between phonology and semantics.

Finally, both CLSM analyses of the imageability advantage identified connections with the occipital cortex, including a parcel-level significant node in the LOC. Neural coding of perceptual features is theorized to be a contributing factor to the imageability advantage, and MEG evidence suggests that the occipital cortex activates in response to imageable words during lexical decision, prior to pMTG activation interpreted as lexical access.^22,78^ Thus, disconnections from extrastriate visual cortex may impair perceptual features from contributing to the imageability advantage.

Our results conflict with a previous analysis localizing lesions affecting concrete word reading (as compared with abstract words) to the left inferior frontal gyrus (IFG)^28^, which was interpreted as reflecting a role of semantic control. However, the word sets used in Dickens et al.^28^ critically did not produce a concreteness effect in reading behavior, complicating the interpretation of the lesion results. In contrast, our stimuli did produce an imageability effect. VLSM did not identify any lesions in the IFG, but CLSM of high imageability irregular words identified a left anterior insula – AG connection. Further, the S-P mapping CLSM analysis identified connections between an MTG parcel and the inferior, middle, and superior frontal gyri. While the frontal gyri may be involved in S-P mapping more broadly, they are unlikely to drive the imageability effect in reading.

While our lesion coverage was robust across the MCA distribution, we did not have left ventral occipitotemporal cortex (lvOT) coverage. One of the most prominent findings in the neuroimaging of reading is the involvement of the left vOT.^79^ While the exact role of the vOT is debated^79–81^, a growing body of research suggests that it is involved in orthographic-semantic processing.^82–84^ Despite poor lesion coverage of the vOT proper, our CLSM analysis can identify disconnections between the lvOT and the rest of the brain that are related to the imageability effect. Indeed, all three CLSM analyses (high imageability words, high imageability irregular words, and S-P mapping) identified disconnections between left temporoparietal cortex and a left inferior temporal parcel (MNI coordinates –50, –44, –18), just anterior to the canonical visual word form area^79^, as reducing the relevant behavioral score. Future research should disambiguate if isolated vOT lesions are sufficient to cause deficits of semantic reading.

### Neural correlates of semantics-phonology mappings

S-P mapping predicted performance on imageable words, controlling for nonverbal semantics and semantic control. This finding provides strong evidence that damaged S-P mappings relate to semantic reading impairments in stroke alexia. STS lesions, extending into the MTG and STG, were associated with the S-P mapping score. This cluster overlapped with the lesions affecting accuracy on imageable words in the lpSTS, providing converging evidence for the lpSTS implementing S-P mapping. Overlapping CLSM results for the S-P score and high imageability word reading provide further demonstration that semantic contributions to reading aloud rely on more general neural systems for semantics and phonology,^17,49,59^ converging in the lSTS.

The disconnections identified in the S-P mapping CLSM analysis implicated similar connections as the imageability analyses, primarily implicating semantic and phonological networks. A MTG node implicated both at the parcel level and in eight of the 17 significant disconnections was anterior to the imageability-related lesions, within territory that has previously been associated with impairments to picture naming^85,86^ and speech comprehension.^87^

Unlike in the imageability analyses, several disconnections between the lMTG (the same parcel described above) and the inferior, middle, and superior frontal gyri related to reduced S-P mapping. While the IFG has been implicated in semantic control^65,88^, the semantic control LSM analyses returned null results. The connections in the S-P analysis mapped onto both posterior IFG areas associated with phonology and anterior areas associated with semantics^89–91^ and perhaps are involved in controlled retrieval of representations, rather than suppression of semantic interference as measured by our semantic control task. The left superior frontal gyrus has been implicated in a number of processes, including working memory^92,93^ and executive control.^94^ These processes may be relevant to the auditory word-to-picture matching task, which required participants to maintain a phonological representation of a word while selecting the target from semantic distractors.

### The computational role of the pSTS in reading aloud

The S-P mapping and imageability VLSM results overlapped in the pSTS. Further, a range of temporoparietal disconnections, associated with both semantic and phonological processing, reduced the imageability advantage. These results together indicate that the pSTS is involved in computing phonological wordforms prior to speech production in reading aloud. Given that we controlled for articulatory impairments and overall reading ability by including low imageability word reading accuracy as a covariate in our analyses, the pSTS cannot serve as the singular endpoint for wordform computation prior to production, as such an endpoint would be shared by all words. Instead, the pSTS likely integrates information from semantic, phonological, and possibly orthographic regions to compute a wordform that can then be translated into a motor program for production.

In the context of computational models of the reading process, this interpretation of the role of the pSTS is most analogous to a “hidden” layer connecting semantics and phonology^7^ in the triangle model of reading.^8^ Hidden layers compute intermediate mappings between other representations.^95^ While both the direct orthography-phonology and the indirect orthography-semantics-phonology pathways contribute to the production of every word^96^, these models suggest that a division of labour develops over learning. Words with regular spelling-to-sound correspondences are primarily mapped directly onto phonology^8^ via a orthography-phonology hidden layer. Inconsistent or irregular words instead rely more on semantically-mediated mappings^7^, reflected in an exaggerated imageability effect as we observed here. The more robust semantic representations of highly imageable words specifically allows them to benefit more from semantic activation.^21^ In typical readers, increased activation may or may not be observed for high relative to low imageability words in the pSTS. Rather, semantic influences may alter the representational geometry of the pSTS. A growing body of evidence suggests that multivariate pattern information can identify altered neural processing without mean activation change.^97,98^ In fact, representational similarity analysis has identified task-dependent semantic effects on vOT^83^ processing, similar to the predictions of this account of pSTS function. Future research should elucidate the representational structure of pSTS activation to understand such an effect.

An alternate interpretation of the results, derived from the DRC model, is that the pSTS implements connections between the semantic system and the phonological lexicon.^6,99^ By this account, damage to the pSTS would impair the capacity of semantics to activate the appropriate whole-word phonological form. However, the DRC model argues that semantics does not play an important role in reading aloud, and that all words can be read correctly without semantic contributions.^6^ The DRC model thus produces no principled reason that accuracy on high imageability words should vary with SP mapping ability when low imageability word reading accuracy is accounted for. Our results support the integral role of semantics in reading aloud, and the interpretation of the pSTS as mediating between phonology and semantics.

The interpretation of the pSTS as mapping between semantics and phonology clarifies and extends a previous study finding that regularization errors were associated with damage to the posterior temporal white matter and MTG.^16^ As the stroke survivors in their sample had only mild semantic impairments, and a picture matching VLSM analysis had minimal overlap with the cluster associated with regularization error production, Binder et al.^16^ interpreted their result as reflecting an S-P impairment. However, they did not directly manipulate a semantic variable. Our high imageability irregular VLSM cluster is most comparable to the Binder et al.^16^ finding, as regularization errors can only occur on irregular words. In the high imageability irregular word VLSM, we identify a similar area of posterior temporal white matter to the regularization finding in Binder et al.^16^. Furthermore, the portion of their finding extending into the MTG overlaps with the S-P mapping VLSM cluster found here. Our results strongly suggest that the regularization errors in the Binder et al.^16^ sample occurred due to a disruption of S-P mappings.

### Two types of acquired semantic reading impairments

Our results, taken in the context of the larger literature, support the existence of at least two types of acquired semantic reading impairment. The first is caused by damage or dysfunction in cortex that subserves semantic representation, as in SD, which leads to deficits on semantic tasks and difficulty reading low frequency irregular words, with regularization errors.^11,12,100^

The second type of acquired semantic reading impairment, demonstrated here in the context of left hemisphere stroke, concerns impaired semantic reading without serious impairment of nonverbal semantic processing. Triangle models suggest a candidate mechanism: a post-semantic impairment that disrupts the ability of semantic activation to contribute to phonology.^96^ This type of semantic reading impairment occurs due to lesions of lpSTS or disconnections causing an inability to integrate semantic and phonological information represented elsewhere. Our results suggest that the pSTS is best characterized as an intermediate processing node, similar to the hidden layer(s) connecting semantics and phonology in triangle models.^7,96^ Further research that better characterizes processing in the pSTS, and reading deficits associated with damage to this region, is warranted to confirm this proposed mechanism in reading aloud.

From a clinical perspective, this work demonstrates that semantic reading deficits occur in post-stroke alexia, and that these deficits can be identified without regard to classical alexia syndromes through a loss of the advantage of high imageability on reading. This process-oriented approach to understanding alexic deficits may help to understand the specific barriers to reading faced by stroke survivors and suggest new behaviors and brain regions to target for reading interventions. For example, training SP mappings along with stimulation of pSTS may be an avenue to improve semantic reading ability after stroke.

## Conclusions

By examining lesions and nonreading semantic abilities that relate to loss of the imageability advantage in word reading, we show that the lpSTS, along with a network of interconnected temporoparietal regions including areas commonly implicated in semantics and phonology, subserves mapping between semantic information and phonology in reading aloud. These results clarify the neurobiology of reading aloud and provide the strongest evidence to date for a post-semantic form of acquired semantic reading disorder that can occur after a left hemisphere stroke.

## References

1 Kjellen, E., Laakso, K. & Henriksson, I. Aphasia and literacy-the insider’s perspective. Int J Lang Commun Disord 52, 573–584 (2017). 10.1111/1460-6984.12302

2 Brookshire, C. E., Wilson, J. P., Nadeau, S. E., Rothi, L. J. G. & Kendall, D. L. Frequency, nature, and predictors of alexia in a convenience sample of individuals with chronic aphasia. Aphasiology 28, 1464–1480 (2014).

3 Coslett, H. B. & Turkeltaub, P. E. in Neurobiology of language 791-803 (Academic Press, 2016).

4 Leff, A. P. & Starrfelt, R. Alexia: Diagnosis, treatment, and theory. (2013).

5 Marshall, J. C. & Newcombe, F. Patterns of Paralexia: A Psyholinguistic Approach. Journal of Psycholinguistic Research 2, 175–199 (1973).

6 Coltheart, M., Rastle, K., Perry, C., Langdon, R. & Ziegler, J. DRC: A dual route cascaded model of visual word recognition and reading aloud. Psych Rev 108, 204–256 (2001).

7 Harm, M. W. & Seidenberg, M. S. Computing the meanings of words in reading: Cooperative division of labor between visual and phonological processes. Psych Rev 111, 662–720 (2004). 10.1037/0033-295X.111.3.662

8 Seidenberg, M. S. & McClelland, J. L. A distributed, developmental model of word recognition and naming. Psychological Review 96, 523–568 (1989).

9 Patterson, K. & Lambon Ralph, M. A. Selective Disorders of Reading? Current Opinion in Neurobiology 9, 235–239 (1999).

10 Plaut, D. C., McClelland, J. L., Seidenberg, M. S. & Patterson, K. Understanding normal and impaired word reading: Computational principles in quasi-regular domains. Psychological Review 103, 56–115 (1996).

11 Woollams, A. M., Lambon Ralph, M. A., Plaut, D. C. & Patterson, K. SD-squared: On the association between semantic dementia and surface dyslexia. Psychological Review 114, 316–339 (2007).

12 Coltheart, M., Masterson, J., Byng, S., Prior, M. & Riddoch, J. Surface dyslexia. Quarterly Journal of Experimental Psychology 35, 469–495 (1983).

13 Hodges, J. R., Patterson, K., Oxbury, S. & Funnell, E. Semantic Dementia: Progressive fluent aphasia with temporal lobe atrophy. Brain 115, 1783–1806 (1992).

14 Bozeat, S., Lambon Ralph, M. A., Patterson, K., Gerrard, P. & Hodges, J. R. Non-verbal semantic impairment in semantic dementia. Neuropsychologia 38, 1207–1215 (2000).

15 Coltheart, M., Tree, J. J. & Saunders, S. J. Computational modeling of reading in semantic dementia: comment on Woollams, Lambon Ralph, Plaut, and Patterson (2007). Psychol Rev 117, 256–271; discussion 271-252 (2010). 10.1037/a0015948

16 Binder, J. R. et al. Surface errors without semantic impairment in acquired dyslexia: a voxel-based lesion-symptom mapping study. Brain 139, 1517–1526 (2016). 10.1093/brain/aww029

17 Lambon-Ralph, M. A., Jefferies, E., Patterson, K. & Rogers, T. T. The neural and computational bases of semantic cognition. Nat Rev Neurosci 18, 42–55 (2017). 10.1038/nrn.2016.150

18 Jefferies, E. & Lambon Ralph, M. A. Semantic impairment in stroke aphasia versus semantic dementia: a case-series comparison. Brain 129, 2132–2147 (2006). 10.1093/brain/awl153

19 Strain, E., Patterson, K. & Seidenberg, M. S. Semantic effects in single-word naming. *Journal of Experimental Psychology: Learning*, Memory, and Cognition 21, 1140–1154 (1995). 10.1037/0278-7393.21.5.1140

20 Strain, E. & Herdman, C. M. Imagibility effects in word naming: An individual differences analysis. Canadian Journal of Experimental Psychology 53, 347–359 (1999). 10.1037/h0087322

21 Plaut, D. C. & Shallice, T. Deep dyslexia: A case study of connectionist neuropsychology. Cognitive Neuropsychology 10, 377–500 (1993). 10.1080/02643299308253469

22 Paivio, A. Dual Coding Theory: Retrospect and Current Status. Canadian Journal of Psychology 45, 255–287 (1991).

23 Friedman, R. B. Recovery from deep alexia to phonological alexia: Points on a continuum. Brain and Language 52, 114–128 (1996).

24 Graves, W. W., Desai, R., Humphries, C., Seidenberg, M. S. & Binder, J. R. Neural systems for reading aloud: a multiparametric approach. Cereb Cortex 20, 1799–1815 (2010). 10.1093/cercor/bhp245

25 Binder, J. R., Westbury, C. F., McKiernan, K. A., Possing, E. T. & Medler, D. A. Distinct brain systems of processing conrete and abstract concepts. Journal of Cognitive Neuroscience 17, 905–917 (2005).

26 Westbury, C. F., Cribben, I. & Cummine, J. Imaging Imageability: Behavioral Effects and Neural Correlates of Its Interaction with Affect and Context. Front Hum Neurosci 10, 346 (2016). 10.3389/fnhum.2016.00346

27 Graves, W. W. et al. Anatomy is strategy: Skilled reading differences associated with structural connectivity differences in the reading network. Brain and Language 133, 1–13 (2014). 10.1016/j.bandl.2014.03.005

28 Dickens, J. V. et al. Localization of Phonological and Semantic Contributions to Reading. J Neurosci 39, 5361–5368 (2019). 10.1523/JNEUROSCI.2707-18.2019

29 Stoel-Gammon, C. The Word Complexity Measure: description and application to developmental phonology and disorders. Clin Linguist Phon 24, 271–282 (2010). 10.3109/02699200903581059

30 Roach, A., Schwartz, M. F., Martin, N., Grewal, R. S. & Brecher, A. The Philadelphia Naming Test: Scoring and Rationale. Clinical Aphasiaology 24, 121–133 (1996).

31 Howard, D. & Patterson, K. Pyramids and Palm Trees: A test of semantic access from pictures and words. (Thames Valley Test Company, 1992).

32 Martin, N., Minkina, I., Kohen, F. P. & Kalinyak-Fliszar, M. Assessment of linguistic and verbal short-term memory components of language abilities in aphasia. J Neurolinguistics 48, 199–225 (2018). 10.1016/j.jneuroling.2018.02.006

33 Fama, M. E. et al. Self-reported inner speech relates to phonological retrieval ability in people with aphasia. Conscious Cogn 71, 18–29 (2019). 10.1016/j.concog.2019.03.005

34 McCall, J. et al. Distinguishing semantic control and phonological control and their role in aphasic deficits: A task switching investigation. Neuropsychologia 173, 108302 (2022). 10.1016/j.neuropsychologia.2022.108302

35 Dickens, J. V. et al. Two types of phonological reading impairment in stroke aphasia. Brain Commun 3, fcab194 (2021). 10.1093/braincomms/fcab194

36 Yushkevich, P. A. et al. User-guided 3D active contour segmentation of anatomical structures: significantly improved efficiency and reliability. Neuroimage 31, 1116–1128 (2006). 10.1016/j.neuroimage.2006.01.015

37 Rorden, C., Bonilha, L., Fridriksson, J., Bender, B. & Karnath, H. O. Age-specific CT and MRI templates for spatial normalization. Neuroimage 61, 957–965 (2012). 10.1016/j.neuroimage.2012.03.020

38 Avants, B. B. et al. A reproducible evaluation of ANTs similarity metric performance in brain image registration. Neuroimage 54, 2033–2044 (2011). 10.1016/j.neuroimage.2010.09.025

39 Tournier, J. D. et al. MRtrix3: A fast, flexible and open software framework for medical image processing and visualisation. Neuroimage 202, 116137 (2019). 10.1016/j.neuroimage.2019.116137

40 Jeurissen, B., Tournier, J. D., Dhollander, T., Connelly, A. & Sijbers, J. Multi-tissue constrained spherical deconvolution for improved analysis of multi-shell diffusion MRI data. Neuroimage 103, 411–426 (2014). 10.1016/j.neuroimage.2014.07.061

41 Smith, R. E., Tournier, J. D., Calamante, F. & Connelly, A. Anatomically-constrained tractography: improved diffusion MRI streamlines tractography through effective use of anatomical information. Neuroimage 62, 1924–1938 (2012). 10.1016/j.neuroimage.2012.06.005

42 Smith, R. E., Tournier, J. D., Calamante, F. & Connelly, A. SIFT2: Enabling dense quantitative assessment of brain white matter connectivity using streamlines tractography. Neuroimage 119, 338–351 (2015). 10.1016/j.neuroimage.2015.06.092

43 Daducci, A. et al. The connectome mapper: an open-source processing pipeline to map connectomes with MRI. PLoS One 7, e48121 (2012). 10.1371/journal.pone.0048121

44 Zhang, Y., Kimberg, D. Y., Coslett, H. B., Schwartz, M. F. & Wang, Z. Multivariate lesion-symptom mapping using support vector regression. Hum Brain Mapp 35, 5861–5876 (2014). 10.1002/hbm.22590

45 DeMarco, A. T. & Turkeltaub, P. E. A multivariate lesion symptom mapping toolbox and examination of lesion-volume biases and correction methods in lesion-symptom mapping. Hum Brain Mapp 39, 4169–4182 (2018). 10.1002/hbm.24289

46 Kimberg, D. Y., Coslett, H. B. & Schwartz, M. F. Power in Vosel-Based Lesion-Symptom Mapping. Journal of Cognitive Neuroscience 19, 1067–1080 (2007).

47 McCall, J. D. et al. Structural disconnection of the posterior medial frontal cortex reduces speech error monitoring. Neuroimage Clin 33, 102934 (2022). 10.1016/j.nicl.2021.102934

48 Mirman, D. et al. Corrections for multiple comparisons in voxel-based lesion-symptom mapping. Neuropsychologia 115, 112–123 (2018). 10.1016/j.neuropsychologia.2017.08.025

49 Binder, J. R., Desai, R. H., Graves, W. W. & Conant, L. L. Where is the semantic system? A critical review and meta-analysis of 120 functional neuroimaging studies. Cereb Cortex 19, 2767–2796 (2009). 10.1093/cercor/bhp055

50 Xia, M., Wang, J. & He, Y. BrainNet Viewer: a network visualization tool for human brain connectomics. PLoS One 8, e68910 (2013). 10.1371/journal.pone.0068910

51 Chang, E. F. et al. Categorical speech representation in human superior temporal gyrus. Nat Neurosci 13, 1428–1432 (2010). 10.1038/nn.2641

52 Wilson, S. M., Bautista, A. & McCarron, A. Convergence of spoken and written language processing in the superior temporal sulcus. Neuroimage 171, 62–74 (2018). 10.1016/j.neuroimage.2017.12.068

53 Indefrey, P. & Levelt, W. J. The spatial and temporal signatures of word production components. Cognition 92, 101–144 (2004). 10.1016/j.cognition.2002.06.001

54 Okada, K. & Hickok, G. Identification of lexical-phonological networks in the superior temporal sulcus using functional magnetic resonance imaging. NeuroReport 17, 1293–1296 (2006).

55 Bouhali, F. et al. Anatomical connections of the visual word form area. J Neurosci 34, 15402–15414 (2014). 10.1523/JNEUROSCI.4918-13.2014

56 Vinckier, F. et al. Hierarchical coding of letter strings in the ventral stream: dissecting the inner organization of the visual word-form system. Neuron 55, 143–156 (2007). 10.1016/j.neuron.2007.05.031

57 Xu, Y., Lin, Q., Han, Z., He, Y. & Bi, Y. Intrinsic functional network architecture of human semantic processing: Modules and hubs. Neuroimage 132, 542–555 (2016). 10.1016/j.neuroimage.2016.03.004

58 Liuzzi, A. G., Aglinskas, A. & Fairhall, S. L. General and feature-based semantic representations in the semantic network. Sci Rep 10, 8931 (2020). 10.1038/s41598-020-65906-0

59 Xu, Y., He, Y. & Bi, Y. A Tri-network Model of Human Semantic Processing. Front Psychol 8, 1538 (2017). 10.3389/fpsyg.2017.01538

60 Boukrina, O. & Graves, W. W. Neural networks underlying contributions from semantics in reading aloud. Front Hum Neurosci 7, 518 (2013). 10.3389/fnhum.2013.00518

61 Dronkers, N. F., Wilkins, D. P., Van Valin, R. D., Jr., Redfern, B. B. & Jaeger, J. J. Lesion analysis of the brain areas involved in language comprehension. Cognition 92, 145–177 (2004). 10.1016/j.cognition.2003.11.002

62 Turken, A. U. & Dronkers, N. F. The neural architecture of the language comprehension network: converging evidence from lesion and connectivity analyses. Front Syst Neurosci 5, 1 (2011). 10.3389/fnsys.2011.00001

63 Hickok, G. P., D. The cortical organization of speech processing. Nature Neuroscience 8, 393–402 (2007).

64 Davey, J. et al. Automatic and Controlled Semantic Retrieval: TMS Reveals Distinct Contributions of Posterior Middle Temporal Gyrus and Angular Gyrus. J Neurosci 35, 15230–15239 (2015). 10.1523/JNEUROSCI.4705-14.2015

65 Noonan, K. A., Jefferies, E., Visser, M. & Lambon Ralph, M. A. Going beyond inferior prefrontal involvement in semantic control: evidence for the additional contribution of dorsal angular gyrus and posterior middle temporal cortex. J Cogn Neurosci 25, 1824–1850 (2013). 10.1162/jocn_a_00442

66 Schwartz, M. F. et al. Anterior temporal involvement in semantic word retrieval: voxel-based lesion-symptom mapping evidence from aphasia. Brain 132, 3411–3427 (2009). 10.1093/brain/awp284

67 Hoffman, P., Lambon Ralph, M. A. & Woollams, A. M. Triangulation of the neurocomputational architecture underpinning reading aloud. Proc Natl Acad Sci U S A 112, E3719–3728 (2015). 10.1073/pnas.1502032112

68 Pobric, G., Jefferies, E. & Ralph, M. A. Amodal semantic representations depend on both anterior temporal lobes: evidence from repetitive transcranial magnetic stimulation. Neuropsychologia 48, 1336–1342 (2010). 10.1016/j.neuropsychologia.2009.12.036

69 Patterson, K., Nestor, P. J. & Rogers, T. T. Where do you know what you know? The representation of semantic knowledge in the human brain. Nat Rev Neurosci 8, 976–987 (2007). 10.1038/nrn2277

70 Woollams, A. M. For richer or poorer? Imageability effects in semantic dementia patients’ reading aloud. Neuropsychologia 76, 254–263 (2015). 10.1016/j.neuropsychologia.2015.03.023

71 Sliwinska, M. W., Khadilkar, M., Campbell-Ratcliffe, J., Quevenco, F. & Devlin, J. T. Early and sustained supramarginal gyrus contributions to phonological processing. Front Psychol 3, 161 (2012). 10.3389/fpsyg.2012.00161

72 Oberhuber, M. et al. Four Functionally Distinct Regions in the Left Supramarginal Gyrus Support Word Processing. Cereb Cortex 26, 4212–4226 (2016). 10.1093/cercor/bhw251

73 Graves, W. W. et al. Correspondence between cognitive and neural representations for phonology, orthography, and semantics in supramarginal compared to angular gyrus. Brain Struct Funct 228, 255–271 (2023). 10.1007/s00429-022-02590-y

74 Damasio, H. & Damasio, A. R. The anatomical basis of conduction aphasia. Brain 103, 337–350 (1980).

75 Pillay, S. B., Stengel, B. C., Humphries, C., Book, D. S. & Binder, J. R. Cerebral localization of impaired phonological retrieval during rhyme judgment. Ann Neurol 76, 738–746 (2014). 10.1002/ana.24266

76 Price, C. J. A review and synthesis of the first 20 years of PET and fMRI studies of heard speech, spoken language and reading. Neuroimage 62, 816–847 (2012). 10.1016/j.neuroimage.2012.04.062

77 Fiez, J. A. & Petersen, S. E. Neuroimaging studies of word reading. Proc Natl Acad Sci U S A 95, 914–921 (1998).

78 Lewis, G. & Poeppel, D. The role of visual representations during the lexical access of spoken words. Brain Lang 134, 1–10 (2014). 10.1016/j.bandl.2014.03.008

79 Cohen, L. et al. The visual word form area: Spatial and temporal characterization of an initial stage of reading in normal subjects and posterior split-brain patients. Brain 123, 291–307 (2000).

80 Price, C. J. & Devlin, J. T. The interactive account of ventral occipitotemporal contributions to reading. Trends Cogn Sci 15, 246–253 (2011). 10.1016/j.tics.2011.04.001

81 Dehaene, S. & Cohen, L. The unique role of the visual word form area in reading. Trends Cogn Sci 15, 254–262 (2011). 10.1016/j.tics.2011.04.003

82 Fischer-Baum, S., Bruggemann, D., Gallego, I. F., Li, D. S. P. & Tamez, E. R. Decoding levels of representation in reading: A representational similarity approach. Cortex 90, 88–102 (2017). 10.1016/j.cortex.2017.02.017

83 Wang, X. et al. Representational similarity analysis reveals task-dependent semantic influence of the visual word form area. Sci Rep 8, 3047 (2018). 10.1038/s41598-018-21062-0

84 Purcell, J. J., Shea, J. & Rapp, B. Beyond the visual word form area: the orthography-semantics interface in spelling and reading. Cogn Neuropsychol 31, 482–510 (2014). 10.1080/02643294.2014.909399

85 Baldo, J. V., Arevalo, A., Patterson, J. P. & Dronkers, N. F. Grey and white matter correlates of picture naming: evidence from a voxel-based lesion analysis of the Boston Naming Test. Cortex 49, 658–667 (2013). 10.1016/j.cortex.2012.03.001

86 Piai, V. & Eikelboom, D. Brain Areas Critical for Picture Naming: A Systematic Review and Meta-Analysis of Lesion-Symptom Mapping Studies. Neurobiol Lang (Camb) 4, 280–296 (2023). 10.1162/nol_a_00097

87 Pillay, S. B., Binder, J. R., Humphries, C., Gross, W. L. & Book, D. S. Lesion localization of speech comprehension deficits in chronic aphasia. Neurology 88, 970–975 (2017).

88 Jackson, R. L. The neural correlates of semantic control revisited. Neuroimage 224, 117444 (2021). 10.1016/j.neuroimage.2020.117444

89 Poldrack, R. A. et al. Functional specialization for semantic and phonological processing in the left inferior prefrontal cortex Neuroimage 10, 15–35 (1999).

90 Yu, M. et al. Neural correlates of semantic and phonological processing revealed by functional connectivity patterns in the language network. Neuropsychologia 121, 47–57 (2018). 10.1016/j.neuropsychologia.2018.10.027

91 Klaus, J. & Hartwigsen, G. Dissociating semantic and phonological contributions of the left inferior frontal gyrus to language production. Hum Brain Mapp 40, 3279–3287 (2019). 10.1002/hbm.24597

92 du Boisgueheneuc, F. et al. Functions of the left superior frontal gyrus in humans: a lesion study. Brain 129, 3315–3328 (2006). 10.1093/brain/awl244

93 Alagapan, S., Lustenberger, C., Hadar, E., Shin, H. W. & Fr hlich, F. Low-frequency direct cortical stimulation of left superior frontal gyrus enhances working memory performance. Neuroimage 184, 697–706 (2019). 10.1016/j.neuroimage.2018.09.064

94 Li, W. et al. Subregions of the human superior frontal gyrus and their connections. Neuroimage 78, 46–58 (2013). 10.1016/j.neuroimage.2013.04.011

95 Hinton, G. E. Connectionist learning procedures. Artificial Intelligence 40, 185–234 (1989).

96 Welbourne, S. R., Woollams, A. M., Crisp, J. & Lambon Ralph, M. A. The role of plasticity-related functional reorganization in the explanation of central dyslexias. Cogn Neuropsychol 28, 65–108 (2011). 10.1080/02643294.2011.621937

97 Kriegeskorte, N., Mur, M. & Bandettini, P. Representational similarity analysis – connecting the branches of systems neuroscience. Front Syst Neurosci 2, 4 (2008). 10.3389/neuro.06.004.2008

98 Kriegeskorte, N. & Wei, X. X. Neural tuning and representational geometry. Nat Rev Neurosci 22, 703–718 (2021). 10.1038/s41583-021-00502-3

99 Perry, C., Ziegler, J. C. & Zorzi, M. Nested incremental modeling in the development of computational theories: the CDP+ model of reading aloud. Psychol Rev 114, 273–315 (2007). 10.1037/0033-295X.114.2.273

100 Patterson, K., Marshall, J. C. & Colthart, M. Surface Dyslexia: Neuropsychological and Cognitive Studies of Phonological Reading. (Lawrence Erlbaum Associates, 1985).

