## Supplemental Materials for "Meaning for reading: The neurocognitive basis of semantic reading impairment after stroke"

**Supplementary Material**

**Materials and methods**

**Participants**

The participants for this study were all a part of an ongoing study on left hemisphere stroke and cognition (R01DC014960). The recruitment period for the present study was from November 2018 to December 2022. People with left hemisphere stroke were recruited from the following sources: author P.E.T.’s outpatient Aphasia Clinic, the Speech and Stroke services at MedStar National Rehabilitation Hospital (NRH), the Stroke National Capital Area Network for Research, the MedStar Georgetown University Hospital Stroke service, regional aphasia and stroke recovery organizations, local stroke support groups, referrals from local clinicians, ClinicalTrials.gov, advertisements in the community, and word of mouth. Neurotypical control participants were recruited from the local community as well as from families of people with left hemisphere stroke. Inclusion criteria included: being a native English speaker (defined as using English as their primary language since age five), having completed 12 or more years of formal education, and having adequate vision and hearing with correction from lenses and hearing aids to complete visual and auditory tests. Education was quantified based on the participant’s highest attained degree in the United States (high school = 12, college = 16, master’s = 18, JD = 19, MD = 20, PhD = 21). Exclusion criteria for both the stroke and control cohorts included: history of neurological disease (including a right hemisphere stroke for the stroke cohort), history of head injury causing loss of consciousness, history of psychiatric disorder requiring hospitalization, ongoing use of psychiatric medications other than common antidepressants, and history of learning disorder requiring educational intervention (e.g., developmental dyslexia). In order to be included in the study, control participants must have scored no less than 1 standard deviation below the mean for their age- and education-matched group on the Montreal Cognitive Assessment.^1^ The participants consisted of all of the controls and stroke survivors from within the recruitment period who were missing none of the relevant behavioral or neuroimaging data.

**Reading assessment**

A list of 200 monosyllabic words varying orthogonally in frequency, spelling-to-sound regularity, and imageability was compiled from published sources.^2–8^ Objective frequency was measured in SUBTLEX-US frequency per million words^9^. The low versus high frequency distinction was defined based on the Zipf scale^10^, with words with Zipf values between 1 and 4 classified as low frequency and words with Zipf values from 4 to 7 (inclusive) classified as high frequency (<https://www.ugent.be/pp/experimentele-psychologie/en/research/documents/subtlexus>). This definition translated to a SUBTLEX-US word frequency of less than 10 per million qualifying as low frequency and a word frequency of at least 10 per million qualifying as high frequency. Our approach to defining regularity followed the two-stage classification system of Strain et al.^3^ Candidate regular and irregular words were first identified based on qualitative adherence to the most frequent grapheme-to-phoneme patterns.^11^ In order to maximize the regular-irregular contrast, we then excluded words based on quantitative spelling-to-sound consistency.^12^ For our exclusionary criteria, we primarily considered consistency at the level of the orthographic body. For example, the target word “tough” can be segmented into the orthographic onset *t–* and the orthographic body *–ough*, which map onto the phonological onset /t/ and rhyme /ʌf/, respectively. The consistencies of the spelling-to-sound mappings were considered for orthographic bodies according to the formula $\frac{\sum{SUBTLEX log}_{10} Word Frequency of Friends}{\sum{SUBTLEX log}_{10} Word Frequency of Friends \& Enemies}$. “Friends” included all monosyllabic words with the same orthographic body and pronunciation, and “enemies” included all monosyllabic words with the same orthographic body but different pronunciation. Completely inconsistent orthographic bodies thus have a value of 0, whereas completely consistent orthographic bodies have a value of 1. All monosyllabic words in the SUBTLEX-US corpus, excluding proper nouns and dialectal variations, were considered in the consistency calculations. Regular words were required to have an orthographic body consistency of 1. For instance, “bike” is a highly consistent regular word in that it only has friends at the level of the body (e.g., “like,” “pike,” “spike”). Irregular words were selected to have maximally inconsistent orthographic bodies. At the level of the body, “tough” is a highly inconsistent irregular word, as it has only one friend, “rough”, and several enemies, such as “through”, “though”, “cough”, and “bough.” Words with inconsistent orthographic onsets were also included. We avoided including categorically irregular words whose onsets or bodies represent highly consistent spelling-to-sound mappings. For example, irregular words beginning with *wa–*, such as “wasp”, or irregular words ending in *–ind*, such as “mind,” were excluded since these categorically irregular grapheme-to-phoneme mappings are highly consistent across similarly spelled words. Irregular words with no orthographic neighbors at the level of the body were included (e.g., “yacht”). In sum, selected regular words have highly consistent grapheme-to-phoneme and body-to-rhyme mappings, and all irregular words have highly inconsistent grapheme-to-phoneme and/or body-to-rhyme mappings. The majority of imageability ratings were drawn from the Cortese & Fugett estimates for 3,000 monosyllabic English words.^13^ Imageability values range from 1 to 7, with words with a rating ≤ 4 classified as having low imageability and words with a rating > 4 classified as having high imageability. Inclusion of function words was minimized in order to avoid confounding part-of-speech effects with the low imageability category. For two words, imageability ratings were not available in the Cortese & Fugett corpus. The imageability for “horse,” a high frequency regular word, was determined by reference to the MRC Psycholinguistic Database.^14^ We estimated the imageability category of “sown,” a low frequency irregular word, by computing the known difference in concreteness^15^ between “sow” and “sown.” First, the 5-point Likert scale concreteness ratings ^15^ were converted to the 7-point Likert scale (1.5x – 0.5). Then, the difference in concreteness between “sow” and “sown” was subtracted from the imageability rating of “sow”.^13^

The word-matching process resulted in a corpus of 200 words that varied orthogonally in frequency, regularity, and imageability (Supplementary tables 1 and 2). High and low frequency words significantly differed in mean frequency per million (t(99.03) = 6.57, *P* < .001) and were matched on mean letter length (t(187.34) = -0.28, p = 0.78), mean imageability rating (t(196.50) = 1.27, *P* = .20), mean articulatory complexity (t(187.86)= -0.71, *P* = .48), and proportion of irregular words. Regular and irregular words were matched on mean letter length (t(193.16) = 0.65, *P* = .51), mean frequency per million (t(189.02) = -0.40, *P* = .69), and mean imageability rating (t(198.00) = 0.03, *P* = .98), but irregular words had lower mean articulatory complexity (t(197.63) = -3.35, *P* < .001). High and low imageability words differed significantly in mean imageability rating t(181.27) = 25.17, *P* < .001), and were matched on mean letter length (t(196.80) = 0.28, *P* = .78), mean frequency per million (t(180.63) = -1.10 *P* = .27), mean articulatory complexity (t(191.99) = -0.31, *P* = .76), and proportion of irregular words.

| Supplementary Table 1. Summary of monosyllabic real word characteristics. | | | | | | |
| --- | --- | --- | --- | --- | --- | --- |
|  | **Frequency** | | **Regularity** | | **Imageability** | |
|  | **High**  (*N* = 100) | **Low**  (*N* = 100) | **Regular**  (*N* = 100) | **Irregular**  (*N* = 100) | **High**  (*N* = 100) | **Low**  (*N* = 100) |
| **Letter Length** |  |  |  |  |  |  |
| Range | 3 – 6 | 3 – 6 | 3 – 6 | 3 – 6 | 3 – 6 | 3 – 6 |
| Mean ± SD | 4.6 ± 0.7 | 4.7 ± 0.8 | 4.6 ± 0.7 | 4.7 ± 0.8 | 4.7 ± 0.7 | 4.6 ± 0.8 |
| **Frequency per million** |  |  |  |  |  |  |
| Range | 10.0 – 921.1 | 0.1 – 9.1 | 0.2 – 921.1 | 0.1 – 828.4 | 0.4 – 840.6 | 0.1 – 921.1 |
| Mean ± SD | 118.2 ± 175.6 | 2.8 ± 2.2 | 64.4 ± 151.2 | 56.7 ± 121.2 | 49.9 ± 113.5 | 71.2 ± 156.4 |
| **Regularity Category** |  |  |  |  |  |  |
| Proportion Irregular | 0.5 | 0.5 | 0 | 1 | 0.5 | 0.5 |
| **Imageability Rating** |  |  |  |  |  |  |
| Range | 1.9 – 6.8 | 1.4 – 6.7 | 1.9 – 6.8 | 1.4 – 6.8 | 4.1 – 6.8 | 1.4 – 4 |
| Mean ± SD | 4.4 ± 1.5 | 4.2 ± 1.4 | 4.3 ± 1.4 | 4.3 ± 1.4 | 5.6 ± 0.8 | 3.0 ± 0.6 |
| **Articulatory Complexity** |  |  |  |  |  |  |
| Range | 0 – 6 | 0 – 7 | 1 – 7 | 0 – 7 | 0 – 7 | 0 – 7 |
| Mean ± SD | 3.1 ± 1.2 | 3.3 ± 1.5 | 3.5 ± 1.4 | 2.9 ± 1.3 | 3.2 ± 1.3 | 3.2 ± 1.5 |

Behavioral testing was conducted at Georgetown University or MedStar National Rehabilitation Hospital in a dedicated testing room using a seventeen-inch Dell laptop with Windows 10 software. The reading assessment was programmed in E-Prime 3.0.^16^ The testing session was video-recorded and participant audio was recorded with a lapel microphone. The laptop was set up in tent-mode and used as a touchscreen. Each trial was cued by a beep, with a word presented at the center of fixation. Participants were instructed to read aloud each word as quickly and accurately as possible. There were three practice trials. Participants advanced to the next trial by pressing an arrow on the bottom left of the screen. Participants had ten seconds to read aloud each word. The stimuli were given in the same pseudorandomized order to all subjects. Accuracy was scored online and rescored offline for the first attempt. The first complete attempt was defined as the first oral response consisting of consonant and vowel, excluding schwa. Isolated vowels were accepted as the first attempt if the target word itself was CV or VC.

Supplementary Table 2. Reading assessment corpus.

| **Word** | **Frequency** | **Regularity** | **Imageability** |
| --- | --- | --- | --- |
| **axe** | Low | Irregular | High |
| **butch** | Low | Irregular | High |
| **choir** | Low | Irregular | High |
| **chrome** | Low | Irregular | High |
| **comb** | Low | Irregular | High |
| **cough** | Low | Irregular | High |
| **crow** | Low | Irregular | High |
| **flood** | Low | Irregular | High |
| **ghoul** | Low | Irregular | High |
| **hearse** | Low | Irregular | High |
| **hearth** | Low | Irregular | High |
| **hoof** | Low | Irregular | High |
| **isle** | Low | Irregular | High |
| **mauve** | Low | Irregular | High |
| **mousse** | Low | Irregular | High |
| **pear** | Low | Irregular | High |
| **pint** | Low | Irregular | High |
| **plaid** | Low | Irregular | High |
| **sew** | Low | Irregular | High |
| **soot** | Low | Irregular | High |
| **suede** | Low | Irregular | High |
| **trough** | Low | Irregular | High |
| **womb** | Low | Irregular | High |
| **wool** | Low | Irregular | High |
| **yacht** | Low | Irregular | High |
| **arc** | Low | Regular | High |
| **arch** | Low | Regular | High |
| **blaze** | Low | Regular | High |
| **claw** | Low | Regular | High |
| **crab** | Low | Regular | High |
| **crate** | Low | Regular | High |
| **crest** | Low | Regular | High |
| **crumb** | Low | Regular | High |
| **flask** | Low | Regular | High |
| **grape** | Low | Regular | High |
| **hedge** | Low | Regular | High |
| **hinge** | Low | Regular | High |
| **hoop** | Low | Regular | High |
| **mulch** | Low | Regular | High |
| **mumps** | Low | Regular | High |
| **pawn** | Low | Regular | High |
| **pike** | Low | Regular | High |
| **plum** | Low | Regular | High |
| **scab** | Low | Regular | High |
| **soil** | Low | Regular | High |
| **spout** | Low | Regular | High |
| **trench** | Low | Regular | High |
| **wedge** | Low | Regular | High |
| **wink** | Low | Regular | High |
| **yield** | Low | Regular | High |
| **ache** | Low | Irregular | Low |
| **awe** | Low | Irregular | Low |
| **bough** | Low | Irregular | Low |
| **brooch** | Low | Irregular | Low |
| **cache** | Low | Irregular | Low |
| **chic** | Low | Irregular | Low |
| **chute** | Low | Irregular | Low |
| **coup** | Low | Irregular | Low |
| **dearth** | Low | Irregular | Low |
| **ewe** | Low | Irregular | Low |
| **fiend** | Low | Irregular | Low |
| **gauge** | Low | Irregular | Low |
| **heir** | Low | Irregular | Low |
| **leapt** | Low | Irregular | Low |
| **phrase** | Low | Irregular | Low |
| **realm** | Low | Irregular | Low |
| **scarce** | Low | Irregular | Low |
| **scheme** | Low | Irregular | Low |
| **seize** | Low | Irregular | Low |
| **sieve** | Low | Irregular | Low |
| **sown** | Low | Irregular | Low |
| **stow** | Low | Irregular | Low |
| **suave** | Low | Irregular | Low |
| **thyme** | Low | Irregular | Low |
| **ton** | Low | Irregular | Low |
| **ail** | Low | Regular | Low |
| **apt** | Low | Regular | Low |
| **breech** | Low | Regular | Low |
| **brisk** | Low | Regular | Low |
| **craze** | Low | Regular | Low |
| **crude** | Low | Regular | Low |
| **dense** | Low | Regular | Low |
| **fame** | Low | Regular | Low |
| **flank** | Low | Regular | Low |
| **gloat** | Low | Regular | Low |
| **hark** | Low | Regular | Low |
| **lurch** | Low | Regular | Low |
| **reap** | Low | Regular | Low |
| **sane** | Low | Regular | Low |
| **scribe** | Low | Regular | Low |
| **sheen** | Low | Regular | Low |
| **sheer** | Low | Regular | Low |
| **slack** | Low | Regular | Low |
| **sledge** | Low | Regular | Low |
| **sole** | Low | Regular | Low |
| **strife** | Low | Regular | Low |
| **surge** | Low | Regular | Low |
| **tame** | Low | Regular | Low |
| **trait** | Low | Regular | Low |
| **yak** | Low | Regular | Low |
| **bear** | High | Irregular | High |
| **blood** | High | Irregular | High |
| **bowl** | High | Irregular | High |
| **break** | High | Irregular | High |
| **bush** | High | Irregular | High |
| **chef** | High | Irregular | High |
| **climb** | High | Irregular | High |
| **death** | High | Irregular | High |
| **dough** | High | Irregular | High |
| **foot** | High | Irregular | High |
| **friend** | High | Irregular | High |
| **heart** | High | Irregular | High |
| **laugh** | High | Irregular | High |
| **push** | High | Irregular | High |
| **shoe** | High | Irregular | High |
| **shove** | High | Irregular | High |
| **son** | High | Irregular | High |
| **steak** | High | Irregular | High |
| **sweat** | High | Irregular | High |
| **sword** | High | Irregular | High |
| **tongue** | High | Irregular | High |
| **touch** | High | Irregular | High |
| **wolf** | High | Irregular | High |
| **wounds** | High | Irregular | High |
| **youth** | High | Irregular | High |
| **beach** | High | Regular | High |
| **belt** | High | Regular | High |
| **bike** | High | Regular | High |
| **black** | High | Regular | High |
| **boat** | High | Regular | High |
| **crane** | High | Regular | High |
| **dance** | High | Regular | High |
| **drank** | High | Regular | High |
| **fire** | High | Regular | High |
| **first** | High | Regular | High |
| **horse** | High | Regular | High |
| **lunch** | High | Regular | High |
| **page** | High | Regular | High |
| **shape** | High | Regular | High |
| **shell** | High | Regular | High |
| **shirt** | High | Regular | High |
| **sit** | High | Regular | High |
| **slept** | High | Regular | High |
| **steam** | High | Regular | High |
| **steel** | High | Regular | High |
| **toast** | High | Regular | High |
| **twelve** | High | Regular | High |
| **wing** | High | Regular | High |
| **witch** | High | Regular | High |
| **yell** | High | Regular | High |
| **blown** | High | Irregular | Low |
| **broad** | High | Irregular | Low |
| **choose** | High | Irregular | Low |
| **deaf** | High | Irregular | Low |
| **doubt** | High | Irregular | Low |
| **flow** | High | Irregular | Low |
| **gross** | High | Irregular | Low |
| **grow** | High | Irregular | Low |
| **heard** | High | Irregular | Low |
| **hour** | High | Irregular | Low |
| **lose** | High | Irregular | Low |
| **most** | High | Irregular | Low |
| **once** | High | Irregular | Low |
| **phase** | High | Irregular | Low |
| **prove** | High | Irregular | Low |
| **put** | High | Irregular | Low |
| **rough** | High | Irregular | Low |
| **soul** | High | Irregular | Low |
| **source** | High | Irregular | Low |
| **swear** | High | Irregular | Low |
| **threat** | High | Irregular | Low |
| **tough** | High | Irregular | Low |
| **tour** | High | Irregular | Low |
| **wear** | High | Irregular | Low |
| **weird** | High | Irregular | Low |
| **bless** | High | Regular | Low |
| **bored** | High | Regular | Low |
| **chance** | High | Regular | Low |
| **dame** | High | Regular | Low |
| **dumb** | High | Regular | Low |
| **fake** | High | Regular | Low |
| **fraud** | High | Regular | Low |
| **grace** | High | Regular | Low |
| **grief** | High | Regular | Low |
| **hate** | High | Regular | Low |
| **help** | High | Regular | Low |
| **late** | High | Regular | Low |
| **must** | High | Regular | Low |
| **past** | High | Regular | Low |
| **proud** | High | Regular | Low |
| **risk** | High | Regular | Low |
| **send** | High | Regular | Low |
| **serve** | High | Regular | Low |
| **stole** | High | Regular | Low |
| **term** | High | Regular | Low |
| **theme** | High | Regular | Low |
| **trust** | High | Regular | Low |
| **week** | High | Regular | Low |
| **wish** | High | Regular | Low |
| **wrote** | High | Regular | Low |

**Neuroimaging**

**Acquisition**

A T1-weighted magnetization prepared rapid gradient echo (MPRAGE) sequence was acquired: 176 sagittal slices; slice thickness = 1 mm; 1 mm^3^ voxels; field of view (FOV) = 256 mm; matrix = 256 x 256 mm; flip angle = 9º; generalized autocalibrating partial parallel acquisition = 2; repetition time (TR) = 1900 ms; echo time (TE) = 2.98 ms; scan time: ~ 5 mins. To quantify white matter connections, multi-shell high angular resolution diffusion imaging (HARDI) scans were acquired: 74 axial slices; slice thickness = 2 mm; diffusion-weighted gradients: 81 directions at b = 3000, 40 at b = 1200, 7 at b = 0; 2 mm^3^ voxels; flip angle = 90º; phase encoding = anterior to posterior; partial Fourier = 6/8; FOV = 232 mm, matrix = 116 x 116 mm; TR = 4700 ms; TE = 82 ms; readout time =  61 ms; slice acceleration = 1; scan time: ~ 10 mins). Six reverse phase-encoded b = 0 images were acquired for susceptibility field estimation (scan time: ~ 1 min). To assist in manual lesion tracing, a fluid-attenuated inversion recovery (FLAIR) sequence was acquired: 192 sagittal slices, slice thickness = 1 mm; 1 mm^3^ voxels; flip angle = 120º; FOV = 256 mm, matrix = 256 mm x 256; TR = 5000 ms; TE = 386 ms; Inversion Time = 1800 ms; scan time: ~ 5 mins.

**Preprocessing**

**Diffusion-weighted images**

Preprocessing of the diffusion-weighted images in MRtrix 3.0^17^ followed the standard stepwise application of Gaussian noise removal, Gibbs ringing artifact removal, correction of distortions induced by motion, eddy currents, and magnetic susceptibility, and inhomogeneity distortion correction. Voxelwise fiber orientation distributions were computed using multi-shell, multi-tissue constrained spherical deconvolution using representative group average response functions for tissue compartments that were derived based on the preprocessed diffusion-weighted images of 25 controls and 25 stroke subjects.^18^ Multi-tissue-informed global intensity normalization was then applied to the fiber orientation distributions generated by spherical deconvolution. A dilated, skull-stripped MPRAGE registered to the mean b = 0 image was supplied as the brain mask for bias field correction, individual response function estimation, and spherical deconvolution. The preprocessed diffusion-weighted images were up-sampled to 1.3 mm prior to spherical deconvolution to increase anatomical contrast for tractography. Structural connectivity was quantified through 15 million streamlines generated by probabilistic anatomically-constrained tractography^19^ on the white matter fiber orientation distributions in native space (algorithm = iFOD2, step = 1, min/max length = 10/300, angle = 45, backtracking allowed, dynamic seeding, streamlines cropped at grey matter-white matter interface). The anatomical image supplied for anatomically-constrained tractography was the five-tissue-type segmented image of the skull-stripped MPRAGE (imputed in stroke subjects).

**Structural image imputation and brain parcellation**

In preparation for structural connectome construction, the structural images of stroke subjects underwent an imputation process prior to processing in FreeSurfer,^20^ which resulted in an image in which the lesioned tissue was filled with normal brain in order to allow tissue segmentation and atlas parcellation. First, the brain was submitted to the ANTS brain extraction script,^21^ including an automatically segmented lesion,^22^ for cost-function masking. This script performed bias field correction and skull-stripping. The extracted brain was then reflected over the x-axis so that the tissue from the spared hemisphere was on the lesioned side, and then registered to the unreflected extracted brain image. As an initial approximation of the expected tissue values, voxels falling within the automatic lesion mask were then filled with values from the reflected image. The resulting image was further repaired via joint image fusion with a reference control group of individuals without lesions. Specifically, a cohort of 25 healthy controls underwent brain extraction and were warped to the lesioned brain in question. The lesioned brain was then sequentially imputed from each of these control brains via ANTS’ joint image fusion algorithm with a search radius of 1. The imputed native space brain was then submitted to FreeSurfer for cortical reconstruction. Lausanne atlas parcellations at scale 125^23^ (<https://github.com/mattcieslak/easy_lausanne>) were generated from the output of Freesurfer’s recon-all for both stroke and control subjects and were registered with the preprocessed diffusion-weighted images via a rigid-body transformation.

Parcel centroid MNI coordinates were derived as follows: first, we warped the native-space MPRAGEs of a library of 39 control subjects to the Clinical Toolbox Older Adult Template^24^ and applied that warp to the native space dilated Lausanne atlas scale 125 images^23^ for those subjects. For each parcel number, voxels in the final combined scale 125 atlas were assigned a parcel value at voxels where more than 90% of subjects had the parcel value. The centroid MNI coordinates were derived as the center of mass of the parcels in this combined image.

**Results**

**Behavioral analyses**

We performed two multiple linear regressions to understand if picture naming (a semantics-to-phonology process) and auditory word to picture matching (a phonology-to-semantics process) were independently related to high- and low-imageability word reading accuracy.

Supplementary Table 3 – Model predicting high imageability word reading accuracy. Significant predictors are bolded.

|  | Accuracy on high imageability word reading | | |
| --- | --- | --- | --- |
| **Predictors** | **Estimates** | **CI** | ***p*** |
| Intercept | -0.219 | [-0.077, 0.514] | 0.143 |
| **Accuracy on low imageability words** | **0.727** | **[0.610, 0.844]** | **<0.001** |
| Nonverbal semantics | 0.015 | [-0.323, 0.352] | 0.340 |
| **Picture Naming** | **0.187** | **[0.054, 0.320]** | **0.007** |
| Auditory Word-to-Picture Matching | -0.174 | [-0.421, 0.072] | 0.162 |
| Semantic control | -0.010 | [-0.093, 0.074] | 0.818 |
| Lesion volume | -0.000 | [-0.000, 0.000] | 0.957 |
| Age | -0.000 | [-0.003, 0.001] | 0.374 |
| Education | 0.006 | [-0.002, 0.000] | 0.072 |
| Observations | 56 |  |  |
| R2 / R2 adjusted | 0.934 / 0.923 |  |  |

Supplementary Table 4 – Model predicting low imageability word reading accuracy. Significant predictors are bolded.

|  | Accuracy on low imageability word reading | | |
| --- | --- | --- | --- |
| **Predictors** | **Estimates** | **CI** | ***p*** |
| Intercept | -0.123 | [-0.077, 0.514] | 0.498 |
| **Accuracy on high imageability words** | **1.057** | **[0.610, 0.844]** | **<0.001** |
| Nonverbal semantics | -0.145 | [-0.323, 0.352] | 0.475 |
| Picture Naming | -0.022 | [0.054, 0.320] | 0.800 |
| Auditory Word-to-Picture Matching | 0.200 | [-0.421, 0.072] | 0.183 |
| Semantic control | 0.010 | [-0.093, 0.074] | 0.835 |
| Lesion volume | -0.000 | [-0.000, 0.000] | 0.245 |
| Age | -0.002 | [-0.003, 0.001] | 0.085 |
| Education | -0.007 | [-0.002, 0.000] | 0.087 |
| Observations | 56 |  |  |
| R2 / R2 adjusted | 0.929 / 0.917 |  |  |

These results suggest that impairment in picture naming, rather than auditory word-to-picture matching, drives reduced accuracy in reading highly imageable words aloud. This is likely because picture naming and oral reading both require the computation of a phonological code from a semantic input, whereas in auditory word-to-picture matching the phonology is provided by the cue (though the participant must maintain that representation).

**Lesion-symptom mapping**

Supplementary Table 5 – VLSM-SVM results for high imageability word reading, high imageability irregular word reading, and S-P mapping.

| **CLSM DV** | **Edge** | **SVR-*β*** | **MNI 1** | **MNI-2** |
| --- | --- | --- | --- | --- |
| All High  Imageability  Word Accuracy | L intraparietal sulcus <> L planum temporale | 9.498 | -24, -73, 26 | -52, -30, 8 |
|  | L middle temporal gyrus <> L dorsal superior temporal sulcus | 9.321 | -60, -22, -16 | -52, -42, 9 |
|  | L posterior inferior temporal gyrus <> L dorsal superior temporal sulcus | 9.145 | -50, -44, -18 | -52, -42, 9 |
|  | L precuneus <> L middle temporal gyrus | 8.838 | -5, -64, 30 | -61, -58, 1 |
|  | L lateral occipital cortex <> L ventral superior temporal sulcus | 8.823 | -17, -96, 19 | -51, -46, 1 |
|  | L angular gyrus <> L lingual gyrus | 8.707 | -35, -74, 44 | -8, -81, -12 |
|  | L supramarginal gyrus <> L middle temporal gyrus | 8.635 | -58, -51, 31 | -60, -22, -16 |
|  | L inferior temporal gyrus <> L dorsal superior temporal sulcus | 8.582 | -47, -11, -39 | -52, -42, 9 |
|  | L cuneus <> L ventral superior temporal sulcus | 8.456 | -10, -87, 37 | -51, -46, 1 |
|  | L middle temporal gyrus <> L dorsal superior temporal sulcus | 8.386 | -61, -39, -9 | -52, -42, 9 |
| High Imageability Irregular Word Accuracy | L intraparietal sulcus <> L planum temporale | 10.000 | -24, -73, 26 | -52, -30, 8 |
|  | L angular gyrus <> L posterior superior temporal gyrus | 9.286 | -58, -51, 31 | -57, -20, -1 |
|  | L supramarginal gyrus <> L angular gyrus | 9.007 | -47, -41, 27 | -39, -56, 23 |
|  | R superior temporal gyrus <> L intraparietal sulcus | 8.905 | 59, -8, -3 | -24, -73, 26 |
|  | L angular gyrus <> L middle temporal gyrus | 8.804 | -44, -58, 41 | -61, -39, -9 |
|  | L angular gyrus <> L anterior superior temporal gyrus | 8.794 | -35, -74, 44 | -51, -8, -9 |
|  | L supramarginal gyrus <> L inferior temporal gyrus | 8.725 | -47, -41, 27 | -50, -26, -26 |
|  | L angular gyrus <> L middle temporal gyrus | 8.686 | -44, -58, 41 | -60, -22, -16 |
|  | L angular gyrus <> L middle temporal gyrus | 8.586 | -58, -51, 31 | -60, -22, -16 |
|  | R thalamus <> L dorsal superior temporal sulcus | 8.575 | 14, -17, 6 | -52, -42, 9 |
|  | L angular gyrus <> L anterior middle temporal gyrus | 8.546 | -58, -51, 31 | -46, 8, -21 |
|  | L angular gyrus <> L inferior temporal gyrus | 8.534 | -58, -51, 31 | -50, -44, -18 |
|  | L anterior insula <> L angular gyrus | 8.427 | -31, 25, 11 | -39, -56, 23 |
|  | L supramarginal gyrus <> L middle temporal gyrus | 8.410 | -47, -41, 27 | -61, -58, 1 |
|  | L lateral occipital cortex <> L ventral superior temporal sulcus | 8.323 | -17, -96, 19 | -51, -46, 1 |
|  | R insula <> L intraparietal sulcus | 8.246 | 35, -19, 10 | -24, -73, 26 |
|  | L supramarginal gyrus <> L middle temporal gyrus | 8.234 | -43, -30, 21 | -61, -58, 1 |
|  | L angular gyrus <> L superior temporal gyrus | 8.217 | -58, -51, 31 | -50, -42, 15 |
|  | L angular gyrus <> L anterior middle temporal gyrus | 8.156 | -44, -58, 41 | -53, -2, -29 |
|  | L angular gyrus <> L lingual gyrus | 8.151 | -58, -51, 31 | -34, -85, -14 |
|  | L superior parietal lobule <> L middle temporal gyrus | 8.007 | -35, -45, 56 | -61, -39, -9 |
|  | L supramarginal gyrus <> L angular gyrus | 7.749 | -47, -41, 27 | -41, -68, 17 |
|  | L lateral occipital cortex* | 4.692 | -43, -75, 4 | - |
|  | L angular gyrus* | 4.316 | -39, -56, 23 | - |
| S-P Score | L superior frontal gyrus <> L middle temporal gyrus | 10.000 | -9, 62, 12 | -60, -22, -16 |
|  | L middle temporal gyrus <> L dorsal superior temporal sulcus | 9.350 | -60, -22, -16 | -52, -42, 9 |
|  | L supramarginal gyrus <> L angular gyrus | 9.310 | -40, -38, 38 | -41, -68, 17 |
|  | L angular gyrus <> L middle temporal gyrus | 9.274 | -44, -58, -9 | -60, -22, -16 |
|  | L postcentral gyrus <> L middle temporal gyrus | 9.223 | -21, -45, 60 | -60, -22, -16 |
|  | L inferior frontal gyrus (p. triangularis) <> L middle temporal gyrus | 9.209 | -41, 39, 15 | -60, -22, -16 |
|  | L calcarine gyrus <> L anterior dorsal superior temporal sulcus | 9.113 | -14, -80, 6 | -57, -20, -1 |
|  | L middle temporal gyrus <> L dorsal superior temporal sulcus | 9.071 | -61, -39, -9 | -52, -42, 9 |
|  | L supramarginal gyrus <> L inferior temporal gyrus | 8.935 | -40, -38, 38 | -50, -26, -26 |
|  | L precentral gyrus <> L middle temporal gyrus | 8.902 | -31, -17, 67 | -60, -22, -16 |
|  | L angular gyrus <> L middle temporal gyrus | 8.896 | -44, -58, -9 | -61, -39, -9 |
|  | L middle frontal gyrus <> L middle temporal gyrus | 8.744 | -33, 53, 1 | -60, -22, -16 |
|  | L orbitofrontal cortex <> L middle temporal gyrus | 8.704 | -31, 25, -11 | -60, -22, -16 |
|  | L supramarginal gyrus <> L anterior superior temporal gyrus | 8.679 | -43, -30, 21 | -46, 8, -21 |
|  | L supramarginal gyrus <> L inferior temporal gyrus | 8.649 | -40, -38, 38 | -50, -44, -18 |
|  | L anterior middle temporal gyrus <> L dorsal superior temporal sulcus | 8.649 | -53, -2, -29 | -52, -42, 9 |
|  | L anterior dorsal superior temporal sulcus <> L putamen | 8.437 | -57, -20, -1 | -23, -3, -2 |
|  | L middle temporal gyrus* | 5.940 | -60, -22, -16 | - |

* = significant at the parcel level (parcel FWE corrected *P* < 0.05), meaning disconnections between the identified parcel and any anatomical endpoint were associated with impaired accuracy.

We performed two VLSM and two CLSM analyses to understand any differences in picture naming (a semantics-to-phonology process) and auditory word to picture matching (a phonology-to-semantics process). Age, education, lesion volume, nonverbal semantics score, semantic control score, and accuracy on low imageability words were regressed out of both analyses. See the Methods for additional details.

Supplementary Table 6 – VLSM-SVM results for picture naming and auditory word to picture matching.

| **Dependent Variable** | **Anatomical Sites** | **Cluster Volume (cc)** | **Center of Mass** | **P-value** |
| --- | --- | --- | --- | --- |
| Picture Naming Accuracy | Middle temporal gyrus  Superior temporal sulcus  Anterior superior temporal gyrus  Inferior temporal gyrus  Temporal white matter | 9.2 | MNI = -56.4, -21.8, -12.6 | 0.01 |
| Auditory Word to Picture Matching Accuracy | Insula  Precentral gyrus  Anterior superior temporal gyrus  Inferior frontal gyrus – pars opercularis  Inferior frontal gyrus – pars triangularis | 7.1 | MNI = -45.9, 0.1, -2.7 | 0.02 |

Supplementary Table 7 – VLSM-SVM results for picture naming and auditory word to picture matching.

| **CLSM DV** | **Edge** | **SVR-*β*** | **MNI 1** | **MNI 2** |
| --- | --- | --- | --- | --- |
| Picture Naming | L postcentral gyrus <> L middle temporal gyrus | 10.000 | -21, -45, 60 | -60, -22, -16 |
|  | L superior frontal gyrus <> L middle temporal gyrus | 9.631 | -9, 62, 12 | -60, -22, -16 |
|  | L supramarginal gyrus <> L inferior temporal gyrus | 9.592 | -40, -38, 38 | -50, -26, -26 |
|  | L angular gyrus <> L middle temporal gyrus | 9.053 | -44, -58, 41 | -60, -22, -16 |
|  | L middle temporal gyrus <> L dorsal superior temporal sulcus | 8.984 | -60, -22, -16 | -52, -42, 9 |
|  | L inferior frontal gyrus (p. orbitalis) <> L middle temporal gyrus | 8.820 | -41, 39, -15 | -60, -22, -16 |
|  | L middle frontal gyrus <> L middle temporal gyrus | 8.598 | -33, 53, 1 | -60, -22, -16 |
|  | L middle temporal gyrus <> L dorsal superior temporal sulcus | 8.548 | -61, -39, -9 | -52, -42, 9 |
|  | L supramarginal gyrus <> L inferior temporal gyrus | 8.539 | -40, -38, 38 | -50, -44, -18 |
|  | L orbitofrontal cortex <> L middle temporal gyrus | 8.500 | -31, 25, -11 | -60, -22, -16 |
|  | L precentral gyrus <> L middle temporal gyrus | 8.446 | -31, -17, 67 | -60, -22, -16 |
|  | L angular gyrus <> L middle temporal gyrus | 8.368 | -44, -58, 41 | -61, -39, -9 |
|  | L supramarginal gyrus <> L angular gyrus | 8.355 | -40, -38, 38 | -41, -68, 17 |
|  | L anterior middle temporal gyrus <> L dorsal superior temporal sulcus | 8.328 | -53, -2, -29 | -52, -42, 9 |
|  | L supramarginal gyrus <> L anterior superior temporal gyrus | 8.308 | -43, -30, 21 | -46, 8, -21 |
|  | L supramarginal gyrus <> L inferior temporal gyrus | 8.301 | -47, -41, 27 | -50, -26, -26 |
|  | L superior parietal lobule <> L middle temporal gyrus | 8.266 | -35, -45, 56 | -60, -22, -16 |
|  | L superior parietal lobule <> L fusiform gyrus | 8.210 | -35, -45, 56 | -50, -56, -10 |
|  | L intraparietal sulcus <> L inferior temporal gyrus | 8.116 | -34, -83, 27 | -50, -44, -18 |
|  | L middle temporal gyrus* | 5.722 | -60, -22, -16 | - |
| Auditory Word to Picture Matching | L inferior frontal gyrus (p. triangularis) <> L medial anterior temporal lobe | 10.000 | -40, 31, 2 | -46, 8, -21 |
|  | L middle frontal gyrus <> L insula | 8.703 | -29, 34, 33 | -37, -5, -2 |
|  | L calcarine gyrus <> L anterior dorsal superior temporal sulcus | 8.699 | -14, -80, 6 | -57, -20, -1 |
|  | L lateral middle frontal gyrus <> L pallidum | 8.699 | -40, 39, 14 | -23, -3, -2 |
|  | L lateral middle frontal gyrus <> L thalamus | 8.680 | -40, 39, 14 | -14, -17, 6 |
|  | L lateral middle frontal gyrus <> L Pallidum | 8.680 | -40, 39, 14 | -19, -4, -5 |
|  | L superior frontal gyrus <> L inferior temporal gyrus | 8.608 | -9, 50, 39 | -47, -11, -39 |
|  | L superior frontal gyrus <> L medial anterior temporal lobe | 8.360 | -9, 50, 39 | -46, 8, -21 |
|  | L inferior frontal gyrus (p. orbitalis) <> L inferior temporal gyrus | 8.301 | -41, 39, -15 | -50, -26, -26 |
|  | L orbitofrontal cortex <> L medial anterior temporal lobe | 8.212 | -31, 25, -11 | -46, 8, -21 |
|  | L medial anterior temporal lobe <> L insula | 8.069 | -46, 8, -21 | -30, 12, 7 |
|  | L superior frontal gyrus <> L insula | 7.932 | -9, 62, 12 | -37, -5, -2 |
|  | L middle frontal gyrus <> L medial anterior temporal lobe | 7.819 | -26, 51, 20 | -46, 8, -21 |
|  | L medial orbitofrontal cortex <> L inferior temporal gyrus | 7.634 | -6, 27, -23 | -47, -11, -39 |
|  | L angular gyrus <> L anterior dorsal superior temporal sulcus | 7.609 | -41, -68, 17 | -57, -20, -1 |
|  | L orbitofrontal cortex <> L ventral anterior temporal lobe | 7.606 | -31, 25, -11 | -34, -16, -32 |
|  | L inferior frontal gyrus (p. opercularis) <> L insula | 7.537 | -47, 11, 11 | -30, 12, 7 |
|  | L inferior frontal gyrus (p. triangularis) <> L anterior middle temporal gyrus | 7.455 | -41, 31, 2 | -53, -2, -29 |
|  | L frontal pole <> L anterior superior temporal gyrus | 7.418 | -8, 66, -10 | -51, -8, -9 |
|  | Rostral middle frontal gyrus <> L anterior superior temporal gyrus | 7.418 | -23, 58, -10 | -51, -8, -9 |
|  | L frontal pole <> L medial anterior temporal lobe | 7.418 | -8, 66, -10 | -46, 8, -21 |
|  | L medial orbitofrontal cortex <> L medial anterior temporal lobe | 7.418 | -8, 50, -10 | -46, 8, -21 |

* = significant at the parcel level (parcel clusterwise *P* < 0.05), meaning disconnections between the identified parcel and any anatomical endpoint were associated with impaired accuracy.

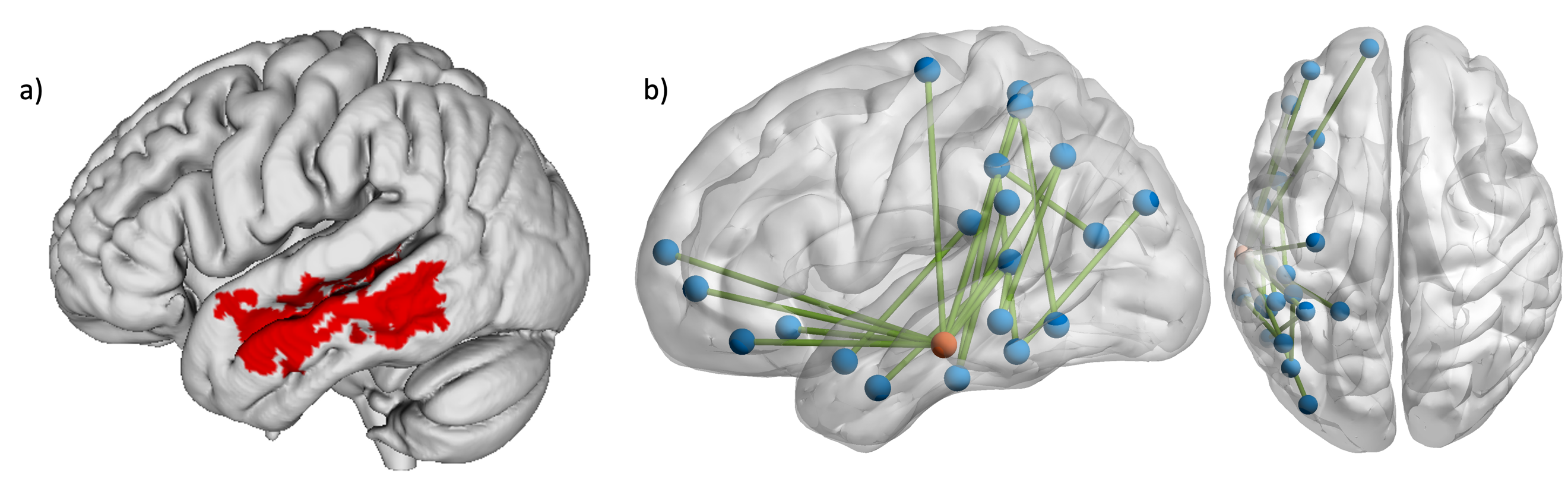

Supplementary Figure 1 – a) SVM-VLSM results (voxelwise *p* < 0.005 and clusterwise FWER *p* < 0.05) and b) SVR-CLSM results (edgewise and parcelwise FWER *p* < 0.05) showing lesion locations and disconnections associated with worse picture naming. Orange node indicates the left middle temporal gyrus node that was significant at the parcel level. Age, education, lesion volume, nonverbal semantics, semantic control, and low imageability word reading accuracy are regressed out of both analyses.

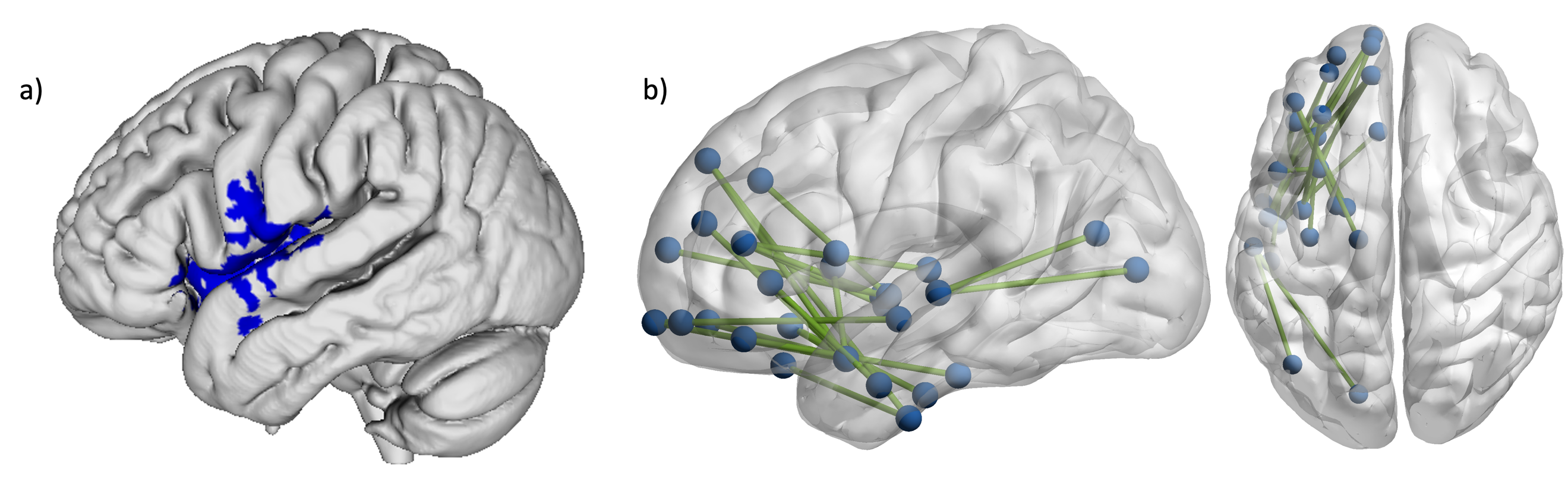

Supplementary Figure 2 – a) SVM-VLSM results (voxelwise *p* < 0.005 and clusterwise FWER *p* < 0.05) and b) SVR-CLSM results (edgewise and parcelwise FWER *p* < 0.05) showing lesion locations and disconnections associated with worse auditory word to picture matching. Age, education, lesion volume, nonverbal semantics, semantic control, and low imageability word reading accuracy are regressed out of both analyses.

The results of these lesion-symptom mapping analyses suggest that the middle temporal gyrus and superior temporal sulcus are essential for picture naming, whereas the anterior portion of the superior temporal gyrus, the precentral gyrus, the insula, and the posterior inferior frontal gyrus (IFG) are needed for auditory word-to-picture matching. Our picture naming VLSM results conform with a previous meta-analyses^26^, and the CLSM results suggest that picture naming draws on a larger network of semantic and phonological cortex. The results for auditory word-to-picture matching are similar to previous findings linking the superior temporal gyrus, insula, posterior IFG, and motor cortex to sensory-motor integration and motor-phonological processing.^27^ In line with the tendency for left-hemisphere stroke to impair phonology more than semantics, these data suggest that the impairment in auditory word-to-picture matching is located in the initial processing of auditory input rather than subsequent semantic processing. CLSM revealed that disconnections within a frontotemporal network impaired auditory word-to-picture matching.

**References**

1.	Rossetti HC, Lacritz LH, Munro Cullum C, Weiner MF. Normative data for the Montreal Cognitive Assessment (MoCA) in a population-based sample. *Neurology*. 2011;77(13):1272-1275. doi:10.1212/WNL.0b013e318230208a

2.	Balota DA, Yap MJ, Hutchison KA, et al. The English Lexicon Project. *Behav Res Methods*. 2007;39(3):445-459. doi:10.3758/BF03193014

3.	Strain E, Patterson K, Seidenberg M. Semantic effects in single word naming. *J Exp Psychol Learn Mem Cogn*. 1995;21(5):1140-1154.

4.	Strain E, Patterson K, Seidenberg MS. Theories of Word Naming Interact with Spelling-Sound Consistency. *J Exp Psychol Learn Mem Cogn*. 2002;28(1):207-214. doi:10.1037/0278-7393.28.1.207

5.	Binder JR, Pillay SB, Humphries CJ, Gross WL, Graves WW, Book DS. Surface errors without semantic impairment in acquired dyslexia: a voxel-based lesion–symptom mapping study. *Brain*. 2016;139:1517-1526.

6.	Behrmann M, Bub D. *Surface Dyslexia and Dysgraphia: Dual Routes, Single Lexicon*. Vol 9.; 1992. doi:10.1080/02643299208252059

7.	Rastle K, Coltheart M. Serial and strategic effects in reading aloud. *J Exp Psychol Hum Percept Perform*. 1999;25(2):482.

8.	Jared D. Spelling-sound consistency and regularity effects in word naming. *J Mem Lang*. 2002;46(4):723-750. doi:10.1006/jmla.2001.2827

9.	Brysbaert M, New B. Moving beyond Kučera and Francis: A critical evaluation of current word frequency norms and the introduction of a new and improved word frequency measure for American English. *Behav Res Methods*. 2009;41(4):977-990. doi:10.3758/BRM.41.4.977

10.	van Heuven WJB, Mandera P, Keuleers E, Brysbaert M. SUBTLEX-UK: A new and improved word frequency database for British English. *Q J Exp Psychol*. 2014;67(6):1176-1190. doi:10.1080/17470218.2013.850521

11.	Venezky RL. *The Structure of English Orthography*. The Hague Mouton Press; 1970.

12.	Jared D. Spelling-sound consistency and regularity effects in word naming. *J Mem Lang*. 2002;46(4):723-750. doi:10.1006/jmla.2001.2827

13.	Cortese MJ, Fugett A. Imageability ratings for 3,000 monosyllabic words. *Behav Res Methods Instrum Comput*. 2004;36(3):384-387.

14.	Coltheart M. The MRC Psycholinguistic Database. *Q J Exp Psychol Sect A*. 1981;33(4):497-505. doi:10.1080/14640748108400805

15.	Brysbaert M, Warriner AB, Kuperman V. Concreteness ratings for 40 thousand generally known English word lemmas. *Behav Res Methods*. 2014;46(3):904-911. doi:10.3758/s13428-013-0403-5

16.	Psychology Software Tools. E-Prime 3.0. Published online 2016.

17.	Tournier JD, Smith R, Raffelt D, et al. MRtrix3: A fast, flexible and open software framework for medical image processing and visualisation. *NeuroImage*. 2019;202:116137. doi:10.1016/j.neuroimage.2019.116137

18.	Jeurissen B, Tournier J donald, Dhollander T, Connelly A, Sijbers J. NeuroImage Multi-tissue constrained spherical deconvolution for improved analysis of multi-shell diffusion MRI data. *NeuroImage*. 2014;103:411-426. doi:10.1016/j.neuroimage.2014.07.061

19.	Smith RE, Tournier JD, Calamante F, Connelly A. Anatomically-constrained tractography: Improved diffusion MRI streamlines tractography through effective use of anatomical information. *NeuroImage*. 2012;62(3):1924-1938. doi:10.1016/j.neuroimage.2012.06.005

20.	Fischl B. NeuroImage FreeSurfer. 2012;62:774-781. doi:10.1016/j.neuroimage.2012.01.021

21.	Avants BB, Tustison NJ, Song G, Cook PA, Klein A, Gee JC. A reproducible evaluation of ANTs similarity metric performance in brain image registration. *NeuroImage*. 2011;54(3):2033-2044. doi:10.1016/j.neuroimage.2010.09.025

22.	Pustina D, Coslett HB, Turkeltaub PE, Tustison N, Schwartz MF, Avants B. Automated Segmentation of Chronic Stroke Lesions Using LINDA : Lesion Identification With Neighborhood Data Analysis. *Hum Brain Mapp*. 2016;37:1405-1421. doi:10.1002/hbm.23110

23.	Daducci A, Gerhard S, Griffa A, et al. The Connectome Mapper: An Open-Source Processing Pipeline to Map Connectomes with MRI. *PLoS ONE*. 2012;7(12). doi:10.1371/journal.pone.0048121

24.	Rorden C, Bonilha L, Fridriksson J, Bender B, Karnath HO. Age-specific CT and MRI templates for spatial normalization. *NeuroImage*. 2012;61(4):957-965. doi:10.1016/j.neuroimage.2012.03.020

25.	Dickens JV, Fama ME, DeMarco AT, Lacey EH, Friedman RB, Turkeltaub PE. Localization of phonological and semantic contributions to reading. *J Neurosci*. 2019;39(27):2707-2718. doi:10.1523/jneurosci.2707-18.2019

26. Piai, V., D. Eikelboom (2023). Brain Areas Critical for Picture Naming: A Systematic

Review and Meta-Analysis of Lesion-Symptom Mapping Studies. *Neurobiol Lang*.

2023;4(2): 280-296.

27. Dickens, J. V. *et al.* Two types of phonological reading impairment in stroke aphasia. *Brain*

*Commun* **3**, fcab194 (2021). <https://doi.org/10.1093/braincomms/fcab194>
